# Type I interferons enhance human dorsal root ganglion nociceptor excitability and induce TRPV1 sensitization

**DOI:** 10.1101/2025.04.22.650068

**Authors:** Úrzula Franco-Enzástiga, Keerthana Natarajan, Felipe Espinosa, Rafael Granja-Vazquez, Hemanth Mydugolam, Theodore J. Price

**Affiliations:** Center for Advanced Pain Studies, School of Behavioral and Brain Sciences, University of Texas at Dallas, Richardson, Texas 75080

**Keywords:** – IFN-α, IFN-β, IFNAR1, IFNAR2, MNK-eIF4E, TRPV1, eFT508

## Abstract

Type I interferons (IFNs) are critical cytokines for antiviral defense and are linked to painful inflammatory diseases like rheumatoid arthritis and neuropathic pain in humans. Studies in rodent models demonstrate a direct action on sensory neurons in the dorsal root ganglion (DRG) to promote hyperexcitability but rodent behavioral results are conflicting with some reports of pro-nociceptive actions and others of anti-nociception. Given the role of type I IFNs in human disease, we sought to clarify the action of action of IFN-α and IFN-β on human DRG (hDRG) nociceptors. We found that IFN receptor subunits IFNAR1 and IFNAR2 are functionally expressed by these neurons and their engagement induces canonical STAT1 signaling and non-canonical MAPK activation as measured by increased phosphorylation of the cap-binding protein eIF4E by MNK1/2 kinases. Using patch clamp electrophysiology, Ca^2+^-imaging, and multi-electrode arrays we demonstrate that IFN-α and -β increase the excitability of hDRG neurons with short (30 min) and long-term (24-48 h) exposure and prolong the duration of capsaicin responses, an effect that is blocked by inhibition of MNK1/2 with eFT508, a specific inhibitor of these kinases. Our studies support the conclusion that type I IFNs are pronociceptive when they interact with hDRG nociceptors in the periphery.

## Introduction

Type I interferons (IFNs) are secreted proteins rapidly induced by viral infection that play a key role in host defense responses (1, 2). They belong to a family of genes contained on chromosome 9 that includes IFN-α, IFN-β, IFN-ε, IFN-κ and IFN-ω in humans. There are 13 genes coding for hIFN-α subtypes that share an amino acid sequence identity ranging from 75% to 99%. Structural differences among type I IFNs account for their varying affinity for the IFN receptor (IFNR) and, consequently, distinct downstream signaling events, such as the phosphorylation of specific signal transducers (3).

Type I IFNs bind to a shared cell surface receptor comprised of 2 transmembrane subunits, IFNAR1 and IFNAR2. After the ternary complex assembly, Janus kinase 1 (JAK1) and Tyrosine kinase 2 (TYK2), which are associated with the cytoplasmic domains of IFNAR1 and IFNAR2, respectively, activate each other by trans-phosphorylation and phosphorylate specific tyrosine residues of IFNAR1 and IFNAR2 (4). Phospho-sites then serve as a docking region for effector proteins of the STAT family. Effector proteins propagate downstream signaling (5). The hallmark of IFN signaling comprises STAT1 and STAT2 phosphorylation, and the formation of a heterotrimer with Interferon Regulatory Factor 9 (IRF9) to form the interferon-stimulated gene factor 3 (ISGF3). ISGF3 translocates to the nucleus to activate the expression of interferon-stimulated genes (ISGs) (6). ISGs functions include inhibition of viral replication by interfering with the viral life cycle, induction of cell death, and activation of innate and adaptative immunity (7). JAKs can induce the formation of STAT1 and STAT3 homodimers that also signal to promote pro-inflammatory responses, or limit them, respectively (8). For antiviral responses, primarily phosphorylation of STAT1/2 is required (5). In addition to JAK-STAT pathway, IFNs can activate non-canonical pathways, such as MAPKs, PI3K/mTOR, and CDKs (9). Notably, while the role of these pathways in IFN signaling is recognized in certain biological systems, their relevance is highly dependent on specific cell types (5).

A pronociceptive effect of IFN-α and IFN-β is supported by experiments in rodents (10–12), by clinical reports of the effects of type I IFN therapy (13), by evidence that ISGs are turned on in the DRG of women with neuropathic pain (14) and by increased type I IFN production in painful diseases like rheumatoid arthritis and lupus (15–17). Guided by the hypothesis that IFNs might induce pain during viral infection, our former work in mice showed that type I IFNs induce nociceptor hyperexcitability and pain when applied to the periphery (11). Furthermore, we showed that they engage mitogen-activated protein (MAP) kinase interacting kinase (MNK) signaling to the elongation initiation factor 4E (eIF4E) pathway (11), a signaling mechanism that is also linked to nociceptor hyperexcitability in humans with neuropathic pain (18, 19). The MNK-eIF4E cascade participates in the translation regulation of specific mRNAs associated with antiviral response, inflammation, metastasis, synaptic plasticity, and pain (20–22). Recently, we demonstrated that Stimulator of Interferon Genes (STING)-IFN-MNK-eIF4E activation underlies pain sensitization caused by a chemotherapeutic agent in mice (12). In the same line, eIF4E has been shown to increase the translation of the E3 ubiquitin ligase tripartite motif protein 32 (TRIM32), enhancing type I IFN signaling and mechanical hypersensitivity in mice (23). Finally, a recent study demonstrated the persistent type I IFN signaling drives MNK-eIF4E-dependent pain in mouse models of rheumatoid arthritis with support for similar signaling in clinical samples from patients (17). Accordingly, inhibiting MNK-mediated phosphorylation of eIF4E has proven beneficial in various preclinical chronic pain models, and painful diseases in humans rendering MNK an important mechanistic target for pharmacological pain relief (11, 17, 18, 21, 22, 24–28).

In contrast to studies suggesting pain induction by IFNs, a recent study showed that CFA-induced inflammation activates STING-IFN-β signaling in nociceptors, leading to decreased nociceptor excitability via upregulation of KChIP1-Kv4.3 (29). Other studies have shown that STING-activated type I IFN signaling reduces neuron excitability by inhibiting the activity of Na^+^ and Ca^2+^ channels (30, 31). In the study of Donnelly and colleagues (30), peripheral administration of IFN-α induced mechanical hypersensitivity that was alleviated by intrathecal administration of IFN-α, suggesting that spinal actions of type I IFNs are anti-nociceptive, an effect consistent with other studies (32–36). Supporting the notion of type I IFNs as antinociceptive agents, a recent work of Defaye and colleagues demonstrates that IFN-β neutralizing antibody administered intrathecally induces thermal hyperalgesia in mice (29). These findings lead to one hypothesis to address these seemingly discordant findings wherein IFNs play a pronociceptive role when applied in the periphery but an antinociceptive role in the CNS. We sought to test if type I IFNs increase the excitability of hDRG neurons.

The goal of our work was to gain mechanistic insight into the action of type I IFNs on human nociceptors. Here, we demonstrate that type I IFNs induce hyperexcitability and sensitization of human nociceptors as well as STAT activation and increased MNK1/2-mediated eIF4E phosphorylation. Furthermore, we provide evidence that one of the pleiotropic effects underlying IFN-induced nociceptor sensitization involves TRPV1 sensitization, an effect blocked by the MNK1/2 inhibitor eFT508. We conclude that type I IFNs sensitize human nociceptors, consistent with the observation that DRG IFN signaling is associated with neuropathic pain in female patients (14) and arthritis pain in animal models and humans (15, 17).

## Results

### IFNAR1 and IFNAR2 mRNAs are expressed in hDRG neurons

Type I IFNs form a ternary complex with the two transmembrane receptor subunits IFNAR1 and IFNAR2 (37, 38). We assessed the mRNA expression of *IFNAR1* and *IFNAR2* in fresh frozen hDRG from organ donors without a history of chronic pain or neuropathy (**Table S1**). In these *in situ* hybridization experiments, each punctum was considered to represent one mRNA of the target. *IFNAR1* was abundantly expressed in all hDRG neurons (**Fig. 1A**). *SCN10A* (+) neurons showed a size-dependent increase in the number of *IFNAR1* mRNAs, suggesting a similar proportion in every neuron independent of its size, as observed in the mRNA number *vs* area correlation plots, where the r value is similar among all neuronal cell sizes (r = 0.62, 0.72, 0.6, respectively, **Fig. S1 A-C**). *IFNAR2* was also expressed in all neurons but at lower levels than *IFNAR1*. The mRNA number *vs* area correlation analysis of *IFNAR2* showed a similar trend in small- and medium-sized neurons, and the level decreased in large neurons (r= 0.43, 0.21 and 0.03, respectively, **Fig. S1 D-F**). Because both subunits are required to induce type I IFN signaling in human cells (39), these results suggest that IFN-α and IFN-β would be expected to have higher activity in small- and medium-sized neurons, likely nociceptors, compared with larger neurons (**Fig. S1 G-H**). Both *IFNAR1* and *IFNAR2* were expressed in *SCN10A* (-) neurons, again with a higher proportion of *IFNAR1* compared to *IFNAR2* (**Fig. 1D**). Our results are in line with spatial and single-cell transcriptomics data (40, 41), showing that both *IFNAR1/2* genes are expressed in neurons with higher abundance for *IFNAR1* (**Fig. 1E**), and also consistent with recent data showing a similar pattern for *IFNAR1* and *IFNAR2* mRNA expression in hDRG neurons (17). Both, *IFNAR1* and *IFNAR2* are also expressed in nonneuronal cell types as shown in single cell sequencing experiments in hDRG (40) (**Fig. 1F**) and in *in situ* hybridization experiments with puncta outside neurons (**Fig. 1A**), aligning to recently published data showing its expression in satellite glial cells (17).

**Figure 1.**
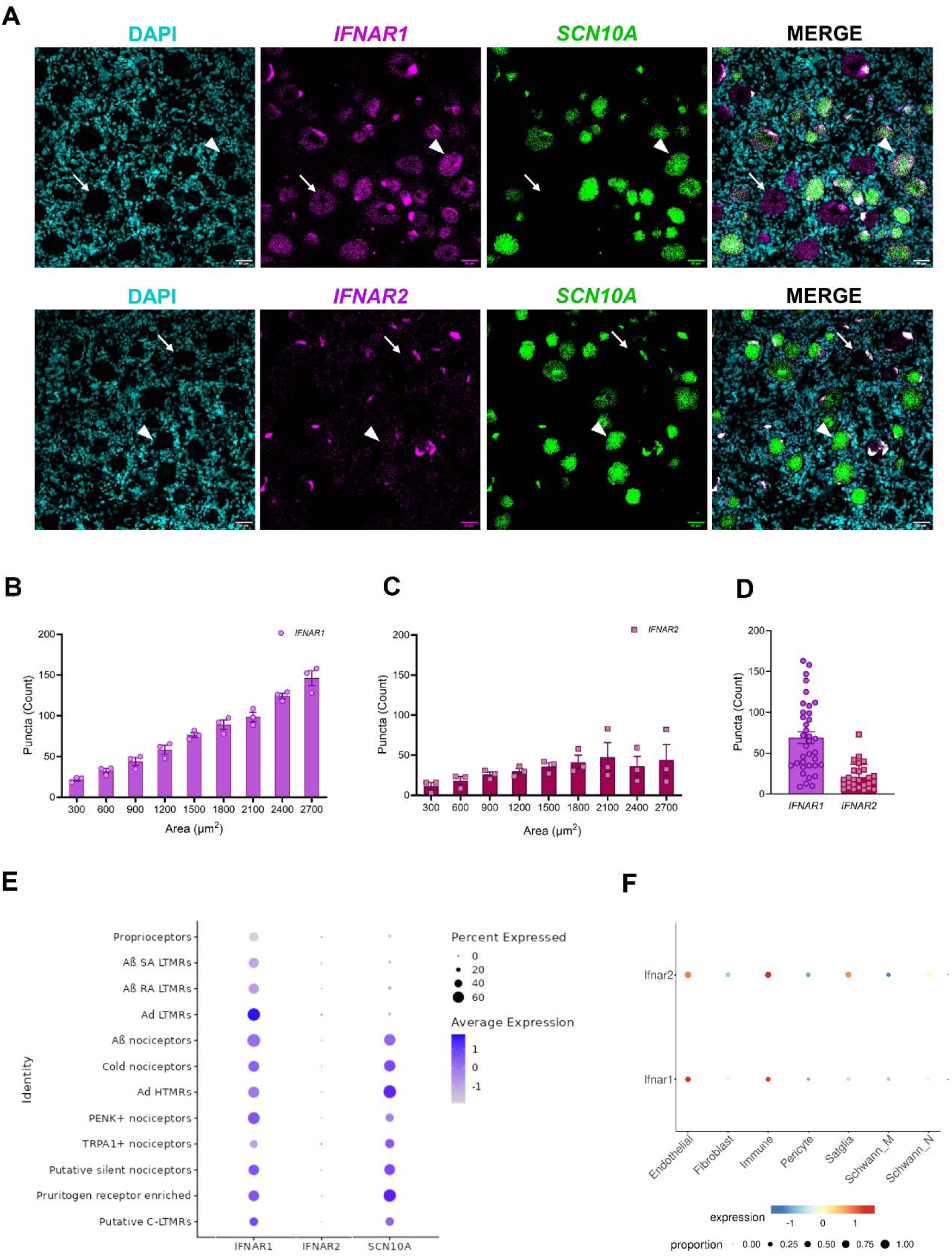
*IFNAR1* and *IFNAR2* mRNAs are expressed in hDRG neurons. **A**. Representative *in situ* hybridization images of *IFNAR1* and *IFNAR2* (magenta) in *SCN10A*(+) neurons (green) in hDRG. Neurons negative and positive for *SCN10A* are indicated by an arrow or arrowhead, respectively. **B-C**. Number of mRNA puncta of *IFNAR1 or IFNAR2* and *SCN10A* as a function of cell area. **D**. Number of *IFNAR1* and *IFNAR2* mRNA puncta in *SCN10A*(-) cells across 3 donors in hDRG. In total, 314 and 470 neurons were analyzed for *IFNAR1* and 470 neurons for *IFNAR2*. **E**. Average expression of *IFNAR1/2* in hDRG neurons according to spatial transcriptomic data of hDRG neurons (41). **F**. Average expression of *IFNAR1/2* in hDRG non-neuronal cell types according to the harmonized atlas of hDRG (40). Scale bar: 50 µm.

### IFNAR1 and IFNAR2 proteins are present in hDRG neurons from organ donors

Based on our RNAscope findings, we tested for the presence of IFNAR subunits in hDRG neurons on cultures prepared from organ donors treated with vehicle or IFNs using immunocytochemistry (ICC). We incubated cultures acutely with 500 U/mL of hIFN-α or hIFN-β (∼1-2 ng/mL), concentration based on previous work on mouse DRG cultures and on other cells (11, 42, 43). In control conditions, IFNAR1 was found to be expressed in peripherin-positive hDRG neurons showing signal in the cytoplasm and in proximity to the cell surface (**Fig. 2A-B**). Neither hIFN-α nor hIFN-β modified IFNAR1 abundance at 30 min or 1 h after stimulation in peripherin-positive hDRG neurons (**Fig. 2A-D**) nor its accumulation in the cell surface (**Fig. 2E-H**). Similar to IFNAR1, in control conditions, IFNAR2 (**Fig. 3A-B**) was detected in hDRG nociceptors, that were also positive with peripherin, in the cytoplasm and proximal to the membrane. IFNAR1 and IFNAR2 availability proximal to the surface suggests the cooperativity among IFNAR subunits to form a ternary complex with IFNs (44), rather than mediating signaling via the activation of IFNAR subunits acting independently of each other (39). hIFN-α increased IFNAR2 abundance in hDRG neurons after 1 h of incubation, whereas hIFN-β did not (**Fig. 3A-D**). This is consistent with hIFN-α’s stronger affinity for IFNAR2 (45). These findings are in line with the notion that upon IFN-α binding, IFNAR2 can be internalized and recycled back to the cell surface (46), as suggested by increased IFNAR2 accumulation proximal to the membrane in hDRG nociceptors (**Fig. 3E,G**). Consistent with our *in situ* hybridization findings, IFNAR1 and IFNAR2 were also detected in nonneuronal cell types in ICC experiments in hDRG cultures. Together, our results confirm that both IFNAR1 and IFNAR2 subunits are translated in hDRG neurons, supporting the idea that endogenous type I IFNs can act directly on human nociceptors.

**Figure 2.**
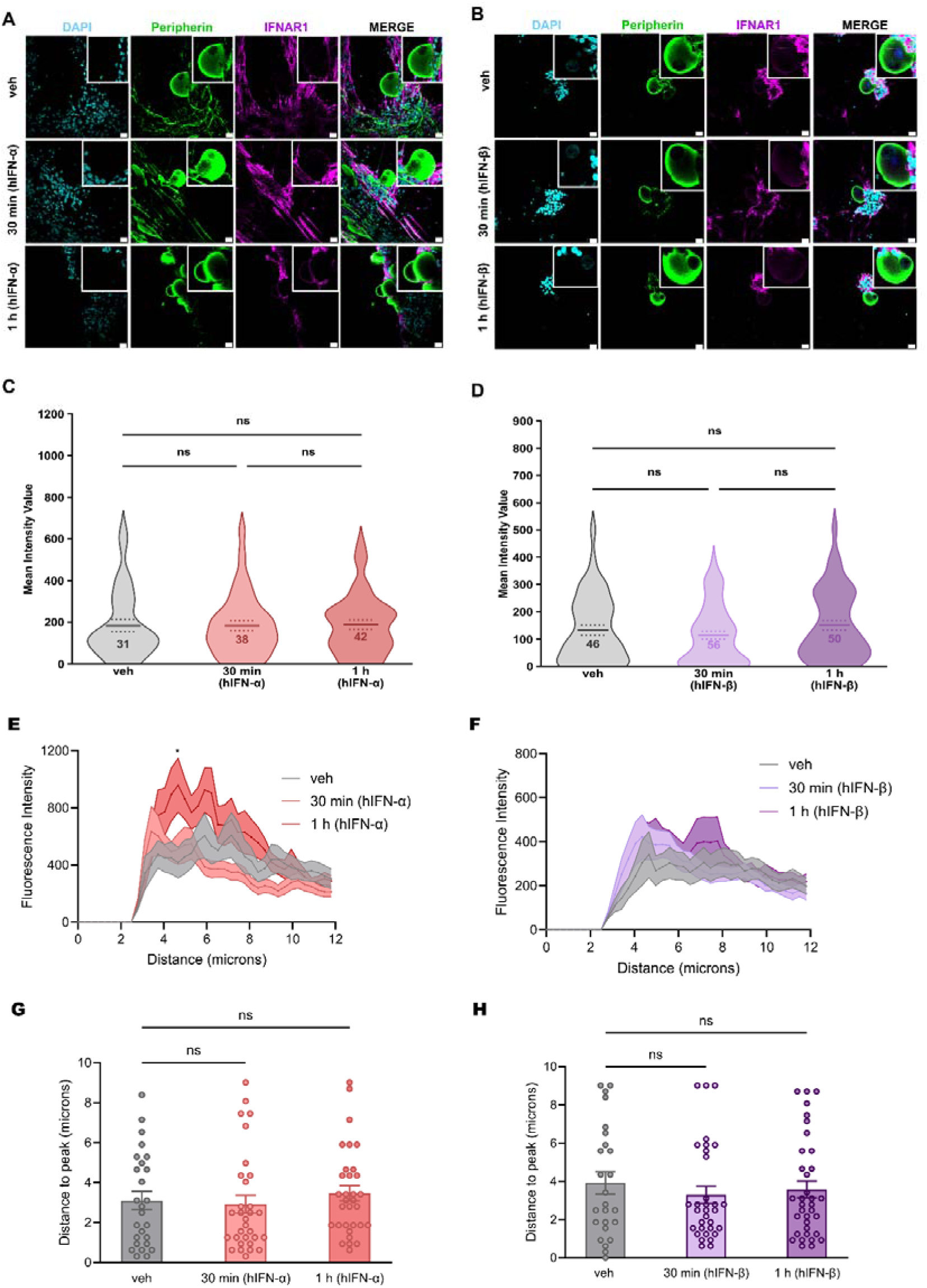
IFNAR1 protein expression in cultured hDRG neuron and response to type I IFN incubation. **A-B.** Representative confocal images of IFNAR1 (magenta) in peripherin positive neurons (green) 30 min or 1 h after 500 U/mL hIFN-α or hIFN-β treatment, respectively. An extra channel was used to subtract lipofuscin autofluorescence signal. **C-D**. Neuronal mean gray intensity value of IFNAR1 in dissociated cultures incubated with hIFN-α or hIFN-β, respectively, for 30 min or 1 h. **E-F**. Mean gray intensity values along perpendicular ROIs hand-drawn to assess IFNAR1 cell surface accumulation signal spanning 10 µm from the exterior of the neuron towards the center of the cell upon hIFN-α or hIFN-β stimulation, respectively. **G-H**. Distance to max gray intensity value for IFNAR1 along the perpendicular ROIs hand-drawn from neurons stimulated with hIFN-α or hIFN-β, respectively. Data are presented as individual values of mean gray value; treatment mean ± SEM are represented, N = 2 organ donors, technical replicates per donor = 2-3 cultures per condition. One-way ANOVA followed by Bonferroni’s test was used to assess group differences in C-D and G-H. *p < 0.05 as determined by two-way ANOVA followed by Bonferroni’s test in E-F. Scale bar: 20 µm. veh: vehicle.

**Figure 3.**
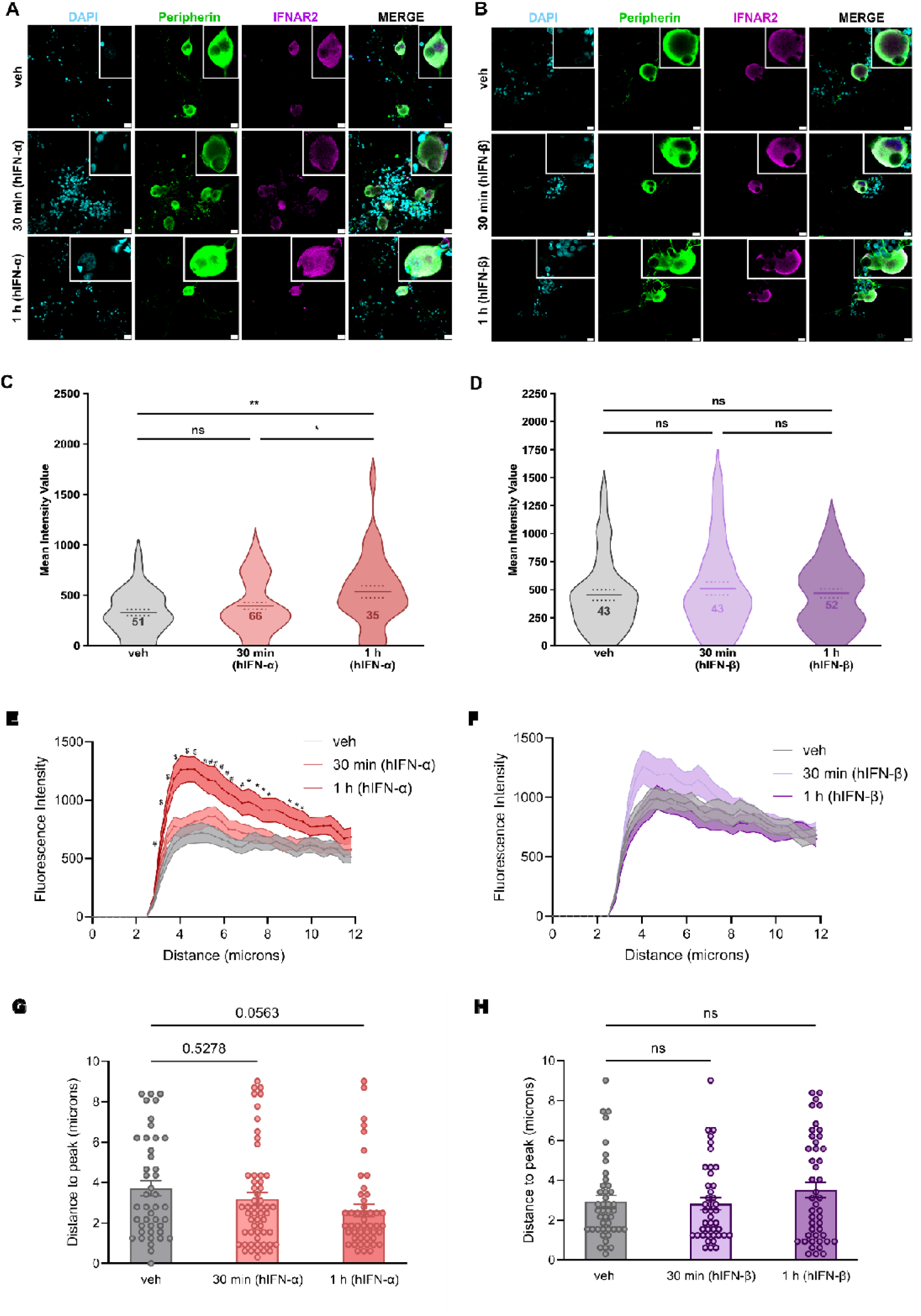
IFNAR2 protein is upregulated by hIFN-α in cultured hDRG neurons. **A**. **A-B**. Representative confocal images of IFNAR2 (magenta) in peripherin positive neurons (green) 30 min or 1 h after 500 U/mL hIFN-α or hIFN-β treatment, respectively. An extra channel was used to subtract lipofuscin autofluorescence signal. **C-D**. Neuronal mean gray intensity value of IFNAR2 in dissociated cultures incubated with hIFN-α or hIFN-β, respectively, for 30 min or 1 h. **E-F**. Mean gray intensity values along perpendicular ROIs hand-drawn to assess IFNAR2 cell surface accumulation signal spanning 10 µm from the exterior of the neuron towards the center of the cell upon hIFN-α or hIFN-β stimulation, respectively. **G-H**. Distance to max gray intensity for IFNAR2 along the perpendicular ROIs hand-drawn from neurons stimulated with hIFN-α or hIFN-β, respectively. Data are presented as individual values of mean gray value; treatment mean ± SEM are represented, N = 2 organ donors, technical replicates per donor = 2-3 cultures per condition. *p < 0.05, **p < 0.01 as determined by one-way ANOVA followed by Bonferroni’s test in C-D and G-H. *p < 0.05, ^#^ p < 0.01, ^$^ p<0.001 as determined by one-way ANOVA followed by Bonferroni’s test in C-D and G-H. *p < 0.05, ^#^ p < 0.01, ^$^ p<0.001 vs veh as determined by two-way ANOVA followed by Bonferroni’s test in E-F. Scale bar: 20 µm. veh: vehicle.

### hIFN-α and hIFN-β induce canonical type I IFN signaling in hDRG neurons

The high abundance of type I IFN receptor mRNA and available protein in hDRG nociceptors and our former IFN-mechanistic work in mouse DRGs (11), led us to explore the activation of type I IFN signaling induced by hIFN-α and hIFN-β application to hDRG cultures. STAT1 is the primary transcription factor mediating cellular response to type I IFNs and is considered a good indicator of IFNR signaling initiation (47). It forms a complex with STAT2, and regulatory factor 9 (IRF9) inducing IFN-stimulated genes (ISGs). Previously, we showed that type I IFN application to cultured DRG neurons in mice induces JAK1 and STAT1 phosphorylation (11). Here, we used ICC to directly test the nuclear accumulation of STAT1 phosphorylation induced by type I IFNs in hDRG neurons labeled with peripherin. hDRG cultures were incubated with type I IFNs (300 and 500 U/mL) for 1 h. We found that hIFN-α and hIFN-β induced a concentration-dependent increase in the phosphorylation of STAT1 (Tyr701) in the nuclear compartment of peripherin positive neurons 1 h after treatment (**Fig. 4A-D**), suggesting early canonical downstream signaling induced by direct stimulation of IFNRs. The magnitude of the increase was two-fold for 500 U/mL hIFN-α and 4.5 times for 500U/mL hIFN-β.

**Figure 4.**
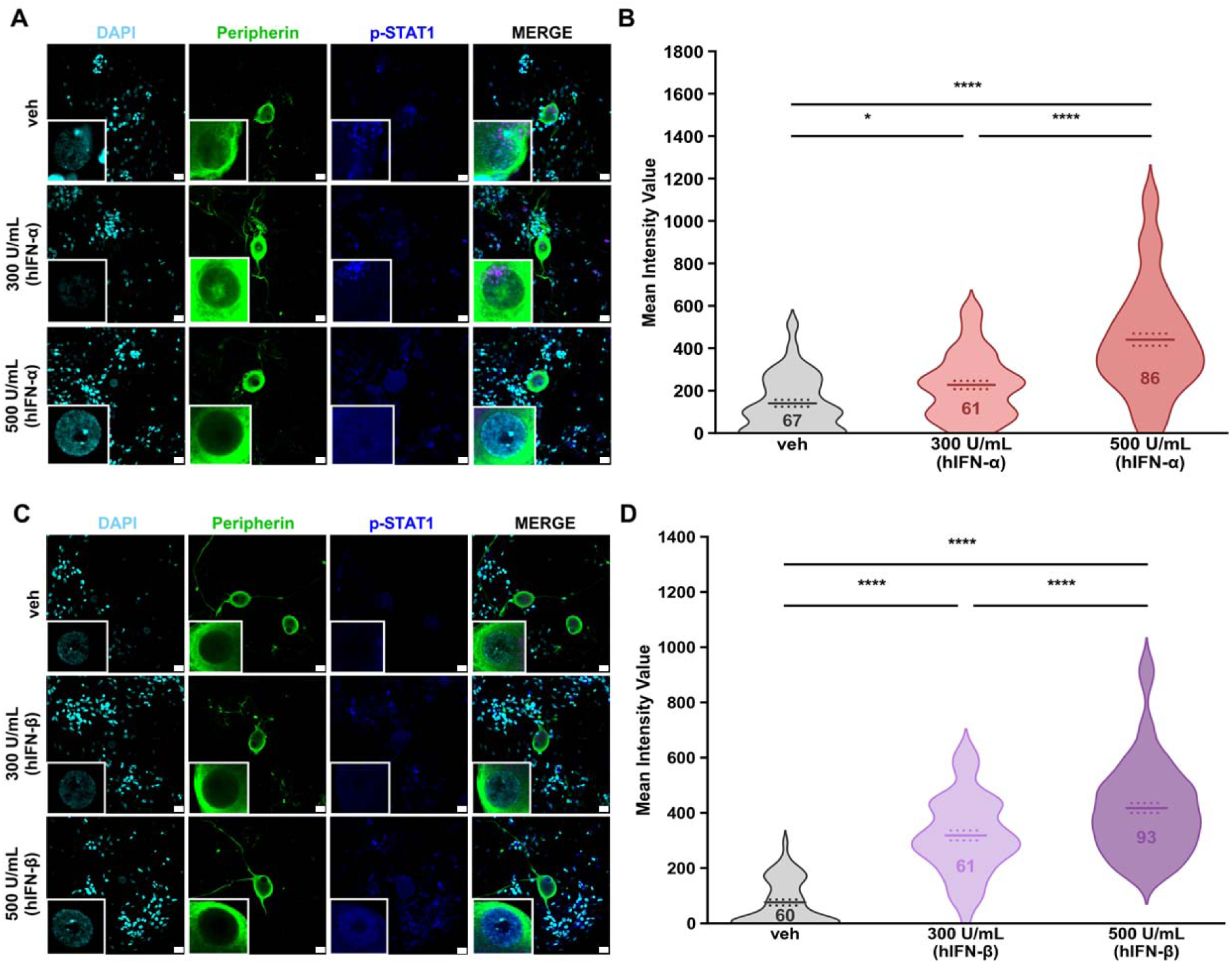
Type I IFN induction of STAT1 signaling in the nuclear compartment of hDRG neurons. **A.** Representative confocal images of p-STAT1 (blue) in peripherin positive neurons (green) 1 h after hIFN-α treatment. An extra channel was used to subtract lipofuscin autofluorescence signal. **B**. Neuronal nuclear mean gray intensity value of p-STAT1 in dissociated cultures incubated with hIFN-α for 1 h. **C**. Representative confocal images of p-STAT1 (blue) in peripherin-positive neurons (green) 1 h after hIFN-β treatment. An extra channel was used to subtract the lipofuscin autofluorescence signal. **D**. Neuronal nuclear mean gray intensity value of p-STAT1 in dissociated cultures incubated with hIFN-β for 1 h. Data are presented as individual values of mean gray value; treatment mean ± SEM are represented, N = 2-3 organ donors, technical replicates per donor = 2-3 cultures per condition. ** p < 0.01, p<0.0001 as determined by one-way ANOVA followed by Bonferroni’s test. Scale bar: 20 µm. veh: vehicle.

### Type I IFN drives human nociceptor excitability after acute and prolonged stimulation

To assess whether the short-term effects of type I IFNs contribute to nociceptor excitability, whole-cell patch-clamp electrophysiological recordings were conducted on cultured hDRG. hDRG nociceptors showed the presence of the hump or shoulder in their action potential waveform characteristic of nociceptors (48, 49). hDRG neuronal cultures were acutely exposed to 500 U/mL of hIFN-α, concentration based on p-STAT1 signaling induction observed in our results. Guided by our RNAscope and ICC experiments showing the expression of type I IFNR in small- and medium-size neurons, electrophysiological recordings were performed on hDRG neurons with these features. Most neurons intended for electrophysiology experiments were covered with glia at dissociation. However, SGCs came off from neurons 1 to 5 days after plating, making it possible to record from cells partially devoid of satellite glial cells (SGCs) (**Fig. 5A**).

**Figure 5.**
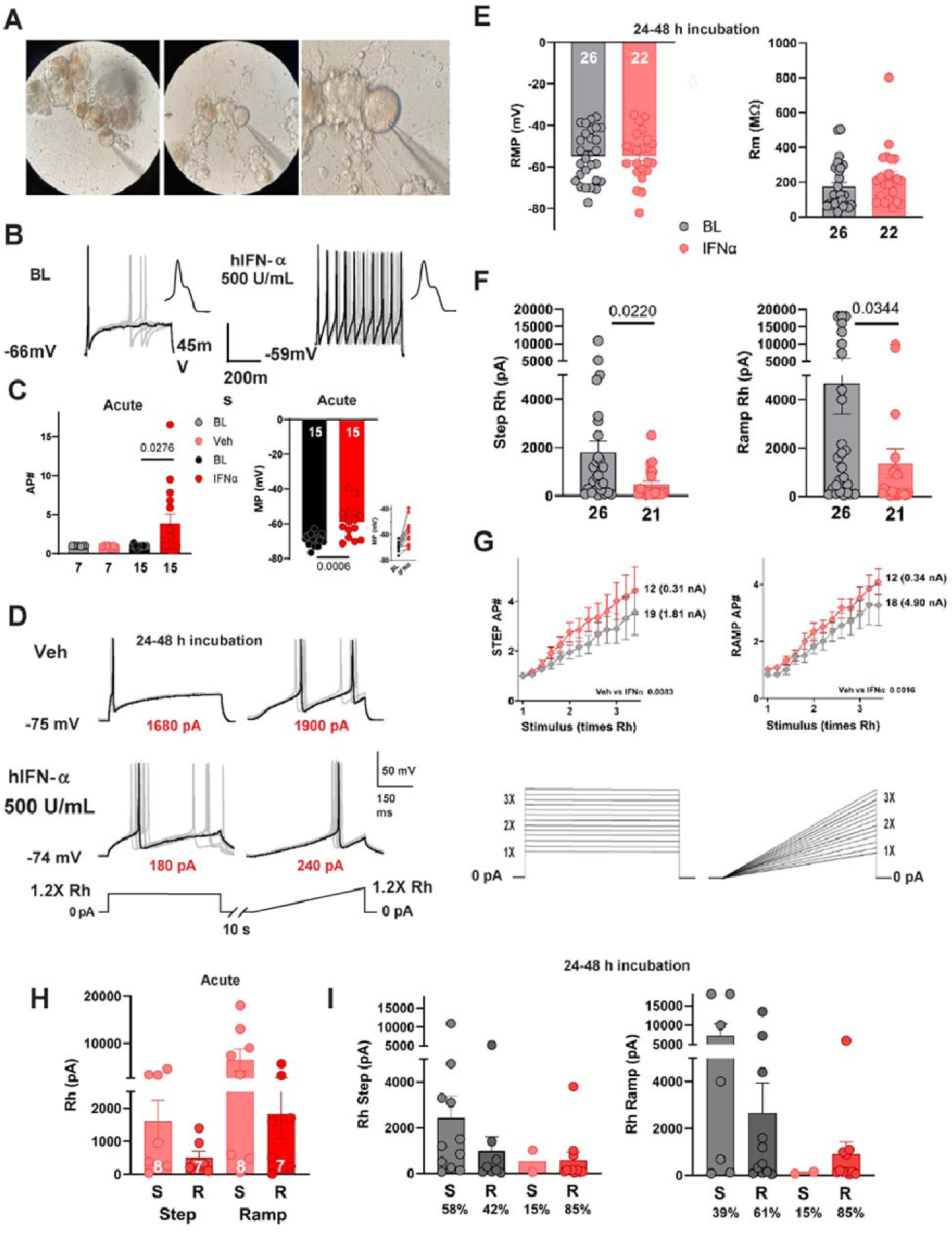
hIFN-α increases the excitability of hDRG neurons. **A.** Pictures of recorded hDRG neurons. The picture on the left shows a hDRG neuron that forms part of a cumulus of cells. The picture in the middle and the zoom on the right show an hDRG neuron where SGCs have migrated to one side after 7 days in culture. **B.** AP firing was induced by step current injections equivalent to 1.2X the rheobase at a holding of -70 mV. Overlapped traces of 10 sweeps before (left) and ∼40 min after start of 500 U/mL hIFN-α perfusion (right). In this cell, the number of APs increased almost 10-fold, accompanied by a mild depolarization of 7 mV. Insets show a zoom of the initial APs in a sweep to highlight the presence of the characteristic AP hump in nociceptors. **C.** Almost half of hDRG neurons responded positively to acute (20-60 min) exposure to hIFN-α. The increase in AP number (#) was significantly different from baseline (BL). In contrast, no difference from baseline (BL) in AP number was found after perfusion of vehicle (left). Acute exposure to hIFN-α also induces a depolarization in hDRG neurons. The inset highlights that most cells were depolarized in the presence of hIFN-α (right). **D.** APs fired upon 1.2X rheobase stimulation after 24-48 h of incubation in either vehicle (top traces) or 500 U/mL of hIFN-α (bottom traces). Numbers in red highlight the difference in rheobase between the experimental groups. **E.** Groups showed no difference in resting membrane potential and in membrane resistance. **F.** After 24-48 h incubation in hIFN-α, rheobase (Rh) was reduced upon both step and ramp stimulation. **G.** Incremental stimulation intensities elicited a higher number of APs after incubation in hIFN-α. Numbers in parenthesis indicate the average rheobase for the corresponding group. **H.** Cells exposed to IFN acutely were separated into IFN responsive or IFN non-responsive groups if, they behaved as single (S, non-responder) or repetitive (R, responsive) spikers after IFN treatment. Rh is shown for step and ramp protocols for S and R cells. **I.** The effect of prolonged exposure to IFN is shown for the percentage of S cells and R cells with (red bars) and without (gray bars) IFN treatment with step and ramp protocols. Data are presented as mean ± SEM. Paired t-test was used to assess group differences in C, and unpaired t-test was used in E and F.

To assess the acute effects of type I IFNs,10 baseline action potentials (APs) elicited at 1.2X rheobase were recorded. This was followed by perfusion of hIFN-α. In 7/15 (47%) of recorded hDRG neurons hIFN-α increased the number of APs within 20-60 min after application (**Fig. 5B-C**). In these neurons, the AP number increased 2-16 fold. In contrast, 0/7 neurons acutely exposed to vehicle showed an increase in AP number. These acute effects are in line with the rapid induction of downstream signaling pathways induced by type I IFNs and with our observations in mouse DRG nociceptors (11). hIFN-α also induced membrane potential depolarization in most cells, with a range of a few millivolts to up to 30 mV (**Fig. 5C**, inset). While some cells that did not respond to hIFN-α with increased AP firing were also depolarized, the depolarization was stronger in cells that did respond to hIFN-α. A significant Pearson correlation coefficient of 0.623 (P = 0.0116) was found when comparing the AP number as a function of the change in membrane potential.

Type I IFNs are known to peak within the first 2 days of viral infection and then there is a decline facilitating the transition to an adaptative immune response characterized by antibody titers (50). However, in some painful disease like lupus there is a persistent increase in type I IFN signaling that can last for years (16), as well as in DRGs from patients with painful rheumatoid arthritis and in the joints of animal models and humans with rheumatoid arthritis (17). We pre-incubated hDRG cultures with 500 U/mL hIFN-α for 24-48 h and performed electrophysiological recordings on human nociceptors. Compared to vehicle treatment, we observed that hIFN-α decreased the step and ramp rheobase (Rh) (**Fig. 5D-G**). Moreover, when exposed to increasing stimulation intensities (see methods), neurons incubated with hIFN-α fired a higher number of APs (**Fig. 5G**). Importantly, because the average rheobase (numbers in parenthesis) was significantly larger in the vehicle group, the difference between groups is more pronounced than they appear in the figure. In contrast to acute experiments, this longer hIFN-α treatment did not affect the membrane potential (**Fig. 5E**). Most DRG neurons responded to intracellular current injections either by firing 1 or 2 APs or with repetitive firing, consistent with what has recently been shown in a patch-seq study of hDRG neurons (51). In our acute experiments, we classified cells as non-responsive to IFN treatment if they did not change their spiking pattern from a single spiker. The rheobase of single spiking cells was higher in both step and ramp current than for repetitive spiking neurons, which are presumptive IFN-responsive cells (**Fig. 5H**). In long-term treatment experiments we observed that IFN shifted the proportion of cells that were repetitive spikers to a higher proportion than without IFN treatment, and also led to an apparent decrease in the rheobase of single spiking neurons (**Fig. 5I**). We did not conduct formal statistical testing on the experiments show in **Fig 5H and I** because of the small sample sizes, but given the recent discovery that repetitive spiking cells have properties of C-fiber nociceptors in hDRG think that this gives important insight into the likely physiology of these cells given recent patch-seq findings (51). Together these results demonstrate that type I IFNs promote hyperexcitability over short (30 – 60 min) and long (24 – 48 h) time scales in hDRG neurons.

### The MNK-eIF4E pathway is activated by type I IFNs in hDRG neurons

Previous work demonstrated that the MNK-eIF4E axis is activated by type I IFNs in mouse DRG neurons and that mechanical nociceptive hypersensitivity caused by type I IFNs in mice are blocked by MNK1/2 knockout or MNK1/2 pharmacological inhibition (11, 17). Using ICC, we tested whether this pathway was induced in hDRG neurons with 300 and 500 U/mL hIFN-α or hIFN-β after 1 h of treatment (**Fig. 6**). Five-hundred U/mL of both IFNs increased the phosphorylation of eIF4E (serine 209), a specific biochemical target of MNK1/2 (52) (**Fig. 6A-D**). Since MNK1/2 activation is associated with increased excitability in human nociceptors (18), this finding suggests that this signaling pathway may underlie effects observed in electrophysiology experiments.

**Figure 6.**
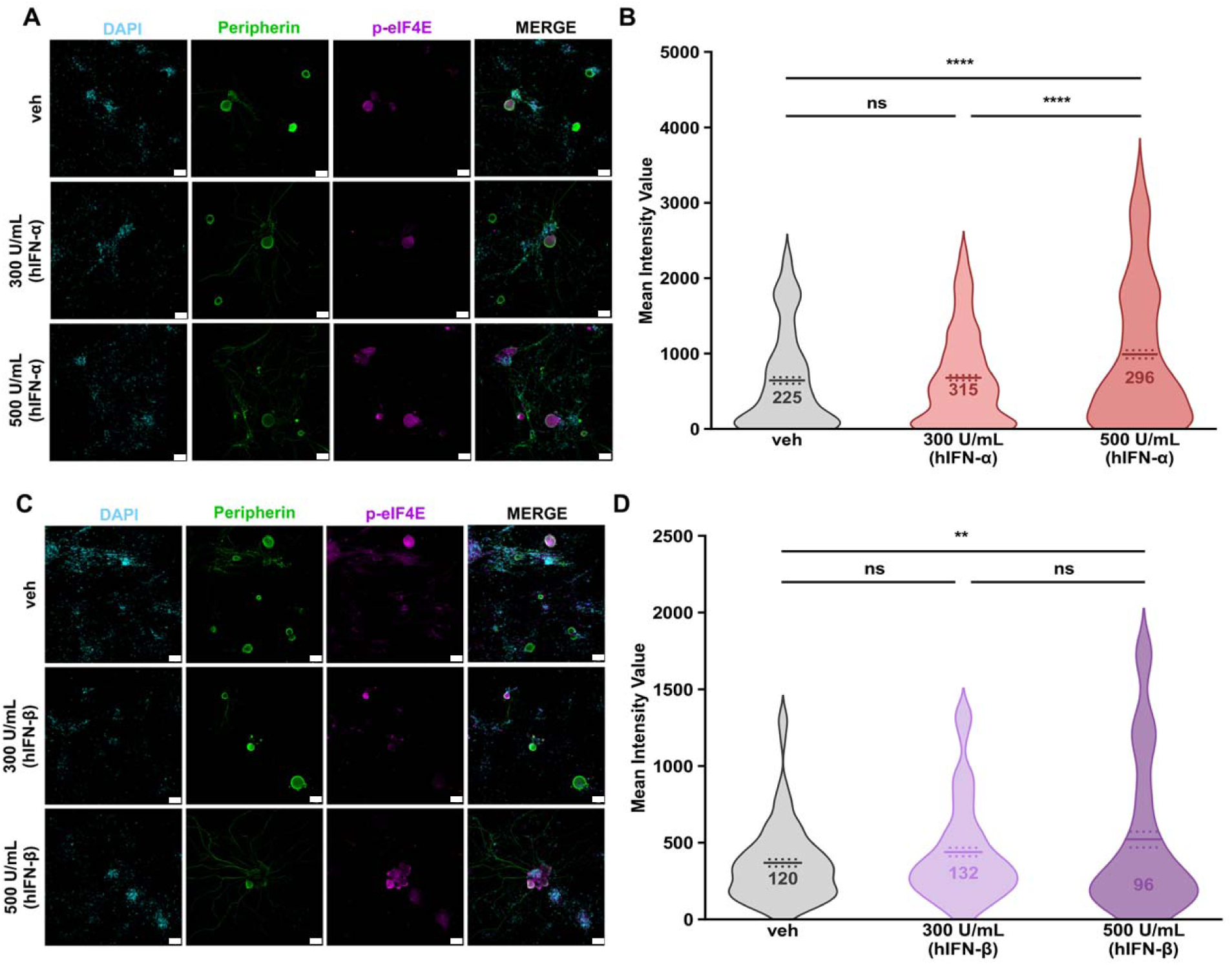
Type I IFNs induce the phosphorylation of eIF4E in hDRG neurons. **A**. Representative images of p-eIF4E (magenta) in peripherin positive neurons (green) 1 h after hIFN-α treatment. **B**. Neuronal mean gray intensity value of p-eIF4E in dissociated cultures incubated with hIFN-α for 1 h **C**. Representative confocal images of p-eIF4E (magenta) in peripherin positive neurons (green) 1 h after hIFN-β treatment. An extra channel was used to subtract the lipofuscin autofluorescence signals in A and C. **D**. Neuronal mean gray intensity value of p-eIF4E in dissociated cultures incubated with hIFN-β for 1 h. Data are presented as individual values of mean gray value; treatment mean ± SEM are represented, N = 2-4 organ donors, technical replicates per donor = 2-3 cultures per condition. **p < 0.01, ****p< 0.0001 as determined by one-way ANOVA followed by Bonferroni’s test. Scale bar: 50 µm. veh: vehicle.

### MNK-eIF4E pathway contributes to sensitization induced by type I IFNs in hDRG neurons

To investigate the implications of the MNK-eIF4E pathway in the IFN-induced effects underlying nociceptor hyperexcitability, we utilized a multiwell micro-electrode array (MEA) platform to complement our patch clamp electrophysiological data. MEA technology has the advantage of allowing the functional monitoring of large neuronal populations and has been successfully utilized to characterize hiPSC-derived nociceptors (53) but has not previously been used with hDRG neurons. APs were recorded in the presence of type I IFNs or vehicle treatment. Independent wells were used to test type I IFNs effects in combination with the specific MNK inhibitor eFT508. APs induced by capsaicin were captured after a 24 h hIFN pre-incubation period with or without eFT508 over a time course of 100 sec after 100 nM capsaicin exposure. Examining TRPV1-mediated responses is relevant because recent studies demonstrate a functional link between type I IFNs and TRPV1 (29, 54, 55). Representative raster plots show increased responsiveness to capsaicin in neurons stimulated with type I IFNs, which is dampened in the presence of eFT508 (**Fig. 7A-B**). Further, an increased AP spike rate in response to capsaicin application was observed with both hIFN-α and hIFN-β, and this effect was abrogated by pre-incubation with eFT508 (**Fig. 7C-D**). The magnitude of the AP spike rate response measured as the area under the curve (AUC) showed a statistically significant increase in the cultures stimulated with hIFNs, an effect that was drastically reduced by the application of eFT508 (**Fig. 7E-F**).

**Figure 7.**
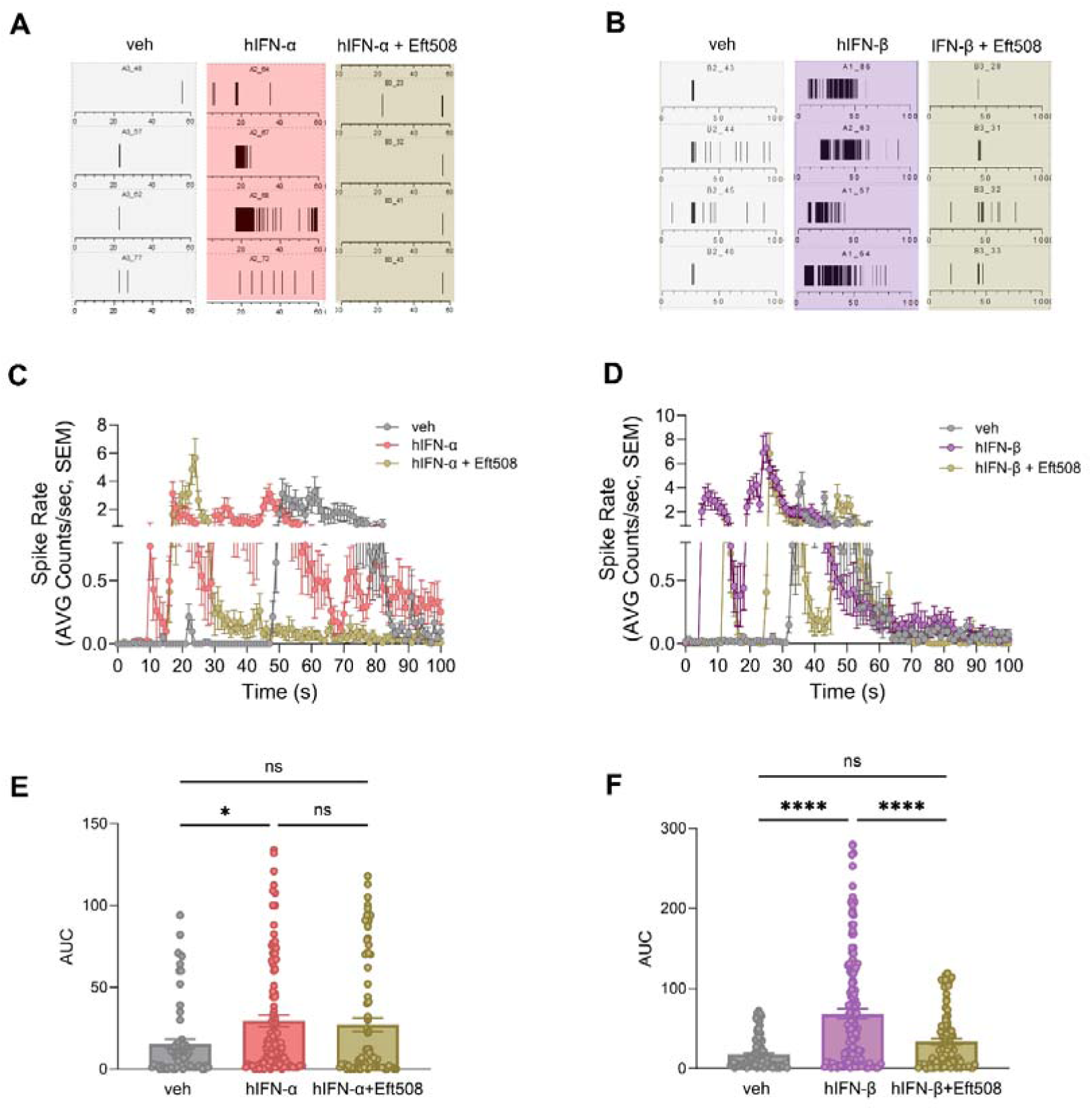
Type I IFNs increase the frequency of APs in hDRG neurons and the MNK inhibitor abrogates this effect. **A-B.** Representative raster plots showing the frequency (spike rate) of APs to capsaicin (100 nM) in neurons pre-treated with vehicle, 500 U/mL hIFN-α or -β, or hIFN-α or -β and 20 nM eFT508 for 24 h. **C.** AP spike rate in response to capsaicin of neurons treated with 500 U/mL hIFN-α or hIFN-α and 20 nM eFT508 for 24 h. **D.** AP spike rate in response to capsaicin of neurons treated with 500 U/mL hIFN-β or hIFN-β and 20 nM eFT508 for 24 h. **E.** AUC as a measure of magnitude of the response in neurons treated with 500 U/mL hIFN-α or hIFN-α and 20 nM eFT508 for 24 h. **F.** AUC as a measure of magnitude of the response in neurons treated with 500 U/mL hIFN-β or hIFN-β and 20 nM eFT508 for 24 h. Data are presented as mean ± SEM. *p<0.05, ****p<0.001 as determined by one-way ANOVA in E-F.

To complement the MEA recordings, Ca^2+^ imaging was used to test nociceptor sensitization via TRPV1 induced by type I IFNs. Pre-incubation with hIFN-α or hIFN-β for 2.5 h was sufficient to induce an increased capsaicin (200 nM) response in hDRG neurons (**Fig. 8A-B**). This statement is based on three observations: 1) the magnitude of the response to capsaicin was significantly increased by both hIFN-α and hIFN-β compared to their respective vehicles as measured by the AUC (**Fig. 8C-D**), 2) the peak Ca^2+^ influx to the cells in response to capsaicin was not statistically changed in presence of hIFN-α (**Fig. 8E**) but was significantly greater with hIFN-β (**Fig. 8F**), and 3) type I IFNs induced a decrease in the latency of the 50% above baseline fluorescence intensity response to capsaicin with hIFN-β treatment, but not hIFN-α (**Fig. 8G-H**). All the cells used in the analysis were small to medium sized neurons, ranging from ∼100 µm^2^ to 800 µm^2^ (**Fig. S2**). These results agree with MEA findings, showing that TRPV1 activity is modulated by type I IFNs in hDRG neurons.

**Figure 8.**
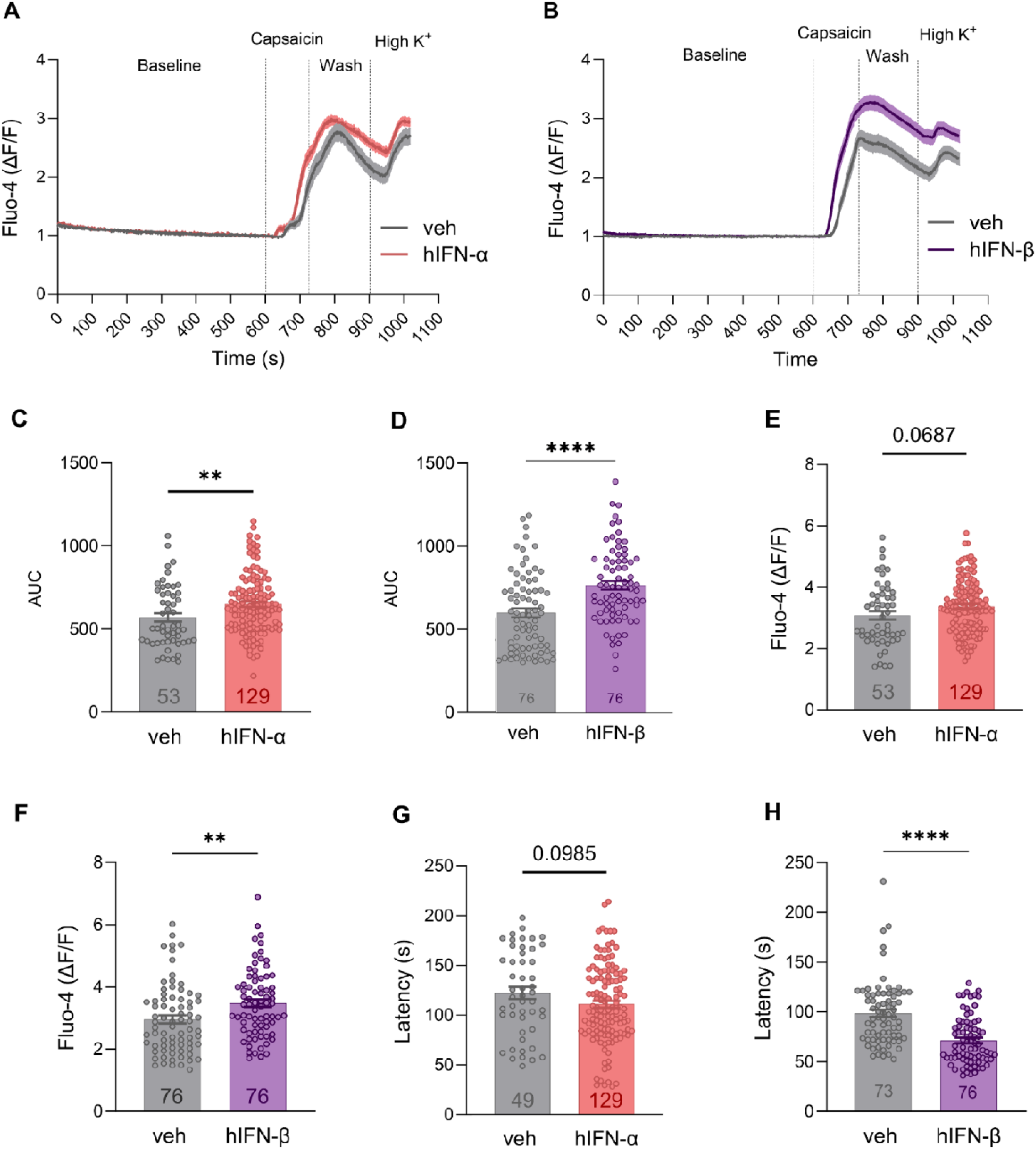
Type I IFN treatment induces sensitization to capsaicin treatment in hDRG nociceptors. **A-B**. Average response traces of hIFN-α and hIFN-β, respectively, or vehicle-treated neurons to capsaicin (200 nM) and high K^+^. **C-D**. Response magnitude to capsaicin of neurons treated with hIFN-α or hIFN-β, respectively, or vehicle. **E, F**. Peak responses to capsaicin in neurons treated with hIFN-α and hIFN-β, respectively, or vehicle. **G, H.** Latency to reach 50% response above baseline to capsaicin in neurons treated with hIFN-α or hIFN-β, respectively, or vehicle. Data are presented as mean ± SEM. **p<0.01, ****p<0.0001 as determined by unpaired t test in C-H.

Together, our findings suggest that type I interferons enhance hDRG nociceptor excitability through a direct activation of IFNAR1/2 engaging MNK-eIF4E pathway and induce TRPV1 sensitization.

## Discussion

The experiments described above provide clear evidence for a sensitizing effect of type I IFNs on hDRG nociceptors. First, we provide evidence for the presence of IFNR subunits on the hDRG cell surface and activation of canonical and non-canonical signaling pathways in hDRG neurons upon activation of these receptors. Second, by performing electrophysiological recordings, we found that acute and prolonged IFN exposure induced hyperexcitability of hDRG neurons, as evidenced by an increase in the number of APs fired in response to step or ramp current depolarization and a decreased rheobase. Third, the IFN-induced sensitized state was also revealed via Ca^2+^ imaging experiments, which showed that type I IFNs induce TRPV1 sensitization. Fourth, in MEA experiments, an increased frequency of APs in response to capsaicin were triggered by type I IFNs, which were abrogated by the specific MNK inhibitor eFT508. Altogether, our findings show that type I IFNs sensitize human nociceptors.

Our initial approach was to determine the expression of IFNAR1 and IFNAR2 in hDRG. IFNAR1/2 participate in the recognition of viral infection like varicella-zoster virus (VZV), herpes virus 1 (HSV-1), HSV-2 (56) and SARS-CoV-2 (57). They also participate in painful inflammatory diseases in humans like rheumatoid arthritis, and neuropathic pain (12, 15, 17, 58). In hDRG neurons, a higher expression of *IFNAR1* mRNA as compared to *IFNAR2* was observed. Imbalance in IFNAR receptor subunit proportion has been observed in different human cell lines and, there is published evidence that the higher abundance of IFNAR1 compensates for the low intrinsic binding of IFN-α to that receptor subunit (46). At the level of protein, both IFNAR1 and IFNAR2 were found in the proximity to the cell surface, suggesting that upon IFN stimulation, the ternary complex is formed (44) to facilitate the transphosphorylation of JAK1 and TYK2 with the subsequent translocation of p-STAT1 to the nucleus, an effect that we also observed in hDRG neurons. p-STAT1, together with p-STAT2, are known to be a primary requisite for the antiviral response of IFNs (5, 47), suggesting that this canonical pathway is induced directly in hDRG nociceptors.

With respect to signaling causing sensitization, we observed that eFT508, a highly potent and specific inhibitor of MNK1 and MNK2 (59), abrogated IFN-sensitization to capsaicin in hDRG neurons. Our former work in mice showed that exogenous application of type I IFN engaged the MNK-eIF4E pathway via MAPK/ERK phosphorylation, inducing tactile allodynia in mice (11). Furthermore, STING activation triggered endogenous type I IFN release in DRGs and sciatic nerves, and induced hypersensitivity in mice, an effect that was reduced in MNK1 KO animals (12). Translation regulation via the MNK-eIF4E axis participates in the induction and maintenance of chronic pain, as shown in different preclinical models (11, 22, 24, 60) and blocking MNK signaling reduces spontaneous activity in nociceptors from humans with neuropathic pain having thoracic vertebrectomy surgery (18). The phosphorylation of p-eIF4E is known to control the translation of mRNAs involved in antiviral response, neuronal plasticity, metastasis, and inflammation (20), actions that have been shown to be therapeutically useful (61). It is also interesting that viruses are known to hijack cellular signaling pathways, and the MNK-eIF4E pathway has been shown to be controlled by viruses such as HSV-1 (62), suggesting an important role of this pathway under infection. In the same vein, it has been shown that one of the effects of activation of the translational pathway MNK-eIF4E is the pronounced upregulation of TRIM32, an E3 ubiquitin ligase that enhances type I IFN signaling and pain (23). Su and colleague’s recent work demonstrated that the underlying cause of joint pain is type I IFN signaling-induced hyperexcitability in Gfra3+ DRG sensory neurons in mice, with a decisive contribution of IFN-activated MNK-eIF4E pathway (11, 12), which aligns well with our findings on human DRG neurons.

We observed that hIFN-β had a more efficacious effect on p-STAT1 induction as compared to hIFN-α, which can be attributed to IFN-β’s higher affinity for IFNAR1 and IFNRA2 (5). The heightened effects for hIFN-β compared to hIFN-α were also observed in calcium imaging and MEA experiments in this work. It’s important to highlight that amino acid sequence identity among hIFN-α and hIFN-β is 30%, a difference that likely underlies their specific functional properties and dissymmetrical affinity for IFNAR1 and IFNAR2 (63). The highest affinity binding hIFN is hIFN-β (5), showing a superior affinity for IFNAR1 (Kd ∼100 nM) and IFNAR2 (Kd ∼0.1 nM) (5), compared with the specific hIFN-α that we used in our experiments, hIFN-α2a (Kd∼ 3.8 µM (64), and ∼1.7 nM, respectively) (45). Another explanation for the heightened hIFN-β effect observed in our study could be that IFN-β complexed with IFNAR subunits has a slower off-rate than IFN-α. IFN-β remains complexed with IFNAR1 for 100 s and with IFNAR2 for 1000 s, while IFN-α has a much faster off-rate of 1 s with IFNAR1 and 100 s with IFNAR2 (5). Our results support the idea that despite sharing a common receptor and inducing similar effects, type I IFNs are not functionally redundant with respect to the sensory system (44), as they show a differential level of biological activity in IFN signaling in hDRG nociceptors.

Functionally, we demonstrate that hIFN-α induces hyperexcitability in hDRG nociceptors. Cells that responded to acute stimulation with hIFN-α showed an increase in the number of APs and membrane potential depolarization indicating hyperexcitability. Prolonged stimulation with hIFN-α enhanced the number of APs and decreased rheobase as well. Interestingly, prolonged IFN stimulation may also induce single spiking cells to behave as repetitive spiker ones. Contradictory findings have been published in rhesus macaque and mouse DRG neurons indicate a fast suppression of AP firing and increased rheobase induced by Universal type I IFN, a hybrid of human IFN-α1 (IFNA1) and IFN-α2a (isoform of IFNA2) (30). This effect was shown to be mediated via sodium and calcium channel activity (30, 31). Interestingly, in this previously published study, APs could not be elicited in hDRG nociceptors, but a hyperpolarization effect was observed. In our study, we analyzed the effects of both type I IFN subtypes and while they did not induce immediate responses they clearly increased nociceptor excitability with short (30 min – 1 h) and prolonged (24 – 48 h) stimulation. Universal type I IFN hybrid differs from human IFN-α2a used for our experiments, as the carboxy-terminal belongs to the sequence of IFN-α1, whereas the NH2-terminal belongs to that of IFN-α2a. These sequence differences can contribute to variations in the biological activity of the hybrid IFN vs the wild-type IFN and may cause the discrepancy between these findings. In line with our current findings and our previous work on mouse DRG (11), another recent study showed that IFN-α (300 U/mL) induces hyperexcitability in mouse nociceptors 1 h after incubation (17). Additionally, neuronal hyperexcitability induced by type I IFNs has been observed in other peripheral ganglia and somatosensory systems. For instance, IFNs directly depolarize the membrane potential of bronchopulmonary afferent neurons from nodose ganglia beyond the action potential threshold (65). IFN-β has also been demonstrated to increase the membrane resistance and the rate of action potential firing in somatosensory neurons of the rat cortex with Ca^2+^-activated K^+^ currents or T-type calcium currents participating in these effects (42). We conclude that our findings are consistent with abundant evidence that type I IFNs increase the excitability of sensory neurons (11, 17) and provide the first evidence that this is the case in human nociceptors.

The work of Defaye and colleagues also differs from the conclusion that type I IFNs sensitize nociceptors. They showed that the activation of STING-IFN pathway by CFA decreases nociceptor excitability through the upregulation of KChIP1-Kv4.3 (29). They also demonstrated a reduction in the number of TRPV1-positive cells in DRGs and decreased TRPV1 expression were observed in mice expressing the human STING mutation N154S in TRPV1 (+) neurons (29). Our MEA and calcium imaging findings contradict those findings and support the conclusion that type I IFNs sensitize human nociceptors. Consistent with our findings, it was recently shown that the direct activation of STING induces Trpv1-dependent sensitization *in vivo*, whereas *in vitro*, it triggered Ca^2+^ responses in capsaicin-responsive neurons, which were lost in Trpv1 KO mice cultures (54). Furthermore, viral DNA induces mechanical hypersensitivity via STING-Trpv1 coupling in mice nociceptors, likely explaining the rapid induction of nocifensive responses to viral ds-DNA in mice (54). Moreover, TRPV1 signaling plays a critical role in the production of antiviral proteins in skin terminals in mice (55), and it has been suggested that proteins like TRPV2 and TRPC1 serve as a port of entry to viruses like HSV-1 into the cell (66). To avoid viral reinfection and spread to bystander cells, TRPV2 is polyubiquitinated by TRIM21 to be degraded in myeloid cells (67). Our results and previous evidence suggest that there is a critical role for the modulation of TRP channels upon viral infection and IFN release. The crosstalk among Trpv1 and type I IFNs is evident in another example. Multiple sclerosis patients using IFN-β as a treatment develop a flu-like syndrome as the most frequent side effect. It presents with muscle pain, fever, malaise, and weakness. Patients carrying an SNP causing a greater maximal TRPV1 channel response (rs222747) showed higher scores of flu-like syndrome symptoms, suggesting that TRPV1 plays a role in pain induction driven by IFN-β (68).

In conclusion, our work shows that type I IFNs increase the excitability of hDRG neurons and engage MNK-eIF4E pathway to induce TRPV1 sensitization. Future experiments are required to study the wide-scale transcriptional changes and protein phosphorylation dynamics induced by type I IFNs in hDRG neurons. Our findings provide substantial insight into states like viral infection and pathologies like neuropathic pain (14), rheumatoid arthritis (15, 17), and lupus (16) that are linked to increased IFN production and pain that can often become chronic and disabling. We propose that in these diseases type I IFNs likely act directly on nociceptors to promote pain and that targeting these cells with therapeutics like MNK inhibitors would be an effective mechanism to reduce pain caused by increased type I IFN signaling.

## Methods

### DRG tissue preparation

Human tissue procurement procedures were approved by the Institutional Review Board at the University of Texas at Dallas. DRGs from human organ donors were obtained through collaboration with the Southwest Transplant Alliance (STA). Right after dissection, lower thoracic or lumbar DRGs from male and female organ donors were either transported in aCSF to be further processed within our facilities or frozen on dry ice right after dissection and stored in a −80 °C freezer as has been described in detail previously (69).

DRG donor demographic information is provided in Table S1. Our inclusion criteria for this study considered organ donors who exhibited no signs of chronic pain or neuropathy and that did not have a history of medications for chronic pain.

### In situ hybridization

hDRGs were embedded in OCT in cryomolds over dry ice and cut into 20 µm sections that were then placed onto SuperFrost® Plus charged slides (Thermo Fisher Scientific, cat. No. 1255015). To allow for the analysis of a diverse population of neurons, three sections separated by at least 100 µm from each other were obtained per donor. Cold 10% formalin (pH 7.4) was used to fix the tissues for 15 min. This was followed by sequential dehydration in 50% ethanol for 5 min, 70% ethanol for 5 min, and 100% ethanol for 10 min at room temperature. The sections were dried briefly and ImmEdge® PAP pen (Vector Laboratories cat. no. H-4000) was used to draw hydrophobic boundaries around the tissue. RNAScope® probes, commercially available from Advanced Cell Diagnostics (ACD), were used to visualize *IFNAR1* (ACD, cat. no. 500891)*, IFNAR2* (ACD, cat. no. 490151), and *SCN10A* (ACD, cat. no. 406291-C2) mRNAs as described previously. In brief, the probes were hybridized at 40 °C for 2 h, followed by amplification. Channel 1 probes were coupled with Cy3 to visualize IFNAR1 and IFNAR2 expression and channel 2 with Cy5 to detect SCN10A. Finally, sections were incubated with 1:5000 DAPI in 1X PBS for 1 min followed by a rinse with 1X PBS before being air-dried completely. ProLong™ Gold Antifade Mountant (Fisher Scientific cat. no. P36930) was used to cover-slip the sections. Sections were imaged at 20X using the Olympus FV3000 RS confocal laser scanning microscope.

The difference in Gaussian (DoG) edge detection method was used to perform analysis on ImageJ Fiji version 2.14.0. ROIs were drawn around each neuron, followed by image duplication of the channel with the target probes. Gaussian Blur with sigma values of 1 and 2 was applied to the duplicated images. The built-in Image Calculator tool was used to subtract images that allowed for puncta detection using the default threshold function. The number of puncta within each ROI was analyzed using the Analyze Particle feature. Data were plotted using Graphpad Prism V9 (Graphpad, San Diego, CA). Neurons with 1 punctum for IFNAR1/2 and 3 puncta for SCN10A were considered positive for the respective targets.

### hDRG cultures

Surgically excised lumbar hDRGs (human dorsal root ganglia) were obtained from male and female organ donors at Southwest Transplant Alliance (STA) approximately 2 h after cross-clamp. We used tissues from both male and female hDRG, as our previous studies showed no sex differences in type I IFN and the IFN-MNK-eIF4E pathways (11). Furthermore, other groups have reported no IFN-signaling differences in DRG neurons although a difference has been observed on microglia; therefore, sex differences at the functional level were not expected and we did not test for them. Right after dissection, the hDRGs were transported in bubbled NMDG-aCSF pH 7.4 (93 mM NMDG, 2.5 mM KCl, 1.25 mM NaH_2_PO_4_, 30 mM NaHCO_3_, 20 mM HEPES, 25 mM glucose, 5 mM ascorbic acid, 2 mM thiourea, 3 mM sodium pyruvate, 10 mM Mg_2_SO_4_, 0.5 mM CaCl_2_, 12 mM N-acetylcysteine; osmolarity 310 mOsm)) on ice to our facilities (70). The hDRGs were minced into small pieces using sterile scissors, immediately transferred to 5 mL of pre-warmed (37°C) STEMxyme 1 (Worthington Biochemical, cat. no. LS004106) solution containing 1 mg/ml STEMxyme in Hank’s Balanced Salt Solution (HBSS), 0.1 mg/mL DNAse I (Worthington Biochemical, cat. no. LS002139), and 10 ng/mL recombinant human β-NGF (R&D Systems, cat. no. 256-GF) and incubated at 37°C in a gently shaking water bath for 8-9 h for ICC and Ca^2+^ imaging. The tissue was triturated using fire-polished sterile glass Pasteur pipettes with decreasing tip diameters until the tissue was completely digested. The cell suspension was passed through a 100-µm cell strainer (VWR, cat. no. 21008-950) and then slowly added to 4 mL of 10% BSA to create a gradient, which was then centrifuged (900 x*g* for 5 min, using a profile of 9 for acceleration and 5 for deceleration) at RT. Neurons were resuspended in BrainPhys® media (Stemcell technologies, cat. no. 05790) supplemented with 1% penicillin/streptomycin, 1% GlutaMAX® (United States Biological, cat. no. 235242), 2% NeuroCult™ SM1 (Stemcell technologies, cat. no. 05711), 1% N-2 Supplement (Thermo Scientific, cat. no. 17502048), 2% HyClone™ Fetal Bovine Serum (ThermoFisher Scientific SH3008803IR), 25 ng/ml recombinant human beta-NGF (R&D Systems, cat. no. 256-GF), 0.15 mg/ml 5-Fluoro-2′-deoxyuridine (Sigma-Aldrich, Cat# F0503-100MG), and 0.35 mg/ml Uridine (Sigma-Aldrich, Cat# U3003-5G). Then 120 µl of the cell suspension was added to a well of an 8-chamber cell culture slides coated with 0.01 mg/mL poly-D-lysine (Sigma, P7405-5MG) and allowed to adhere for 2 h at 37°C and 5% CO_2_. The chambers were then flooded with 0.65 mL of media. The cell cultures were maintained at 37°C with 5% CO_2_ for 5-6 days before treatment. Culture media was changed every other day.

For electrophysiological recordings and MEA experiments, hDRGs were dissociated similarly as above but with slight differences. hDRGs were weighed and 15-25 mg of minced hDRG were added per mL of dissociation solution and incubated at RT on a nutator shaker rotating at 62 rpm. Cells were dissociated into two batches. The first cell batch was obtained following 90-120 min of initial incubation. Fresh dissociation solution was added to the remaining tissue for an additional 30-60 min. At the expense of a lower yield, these shorter periods of dissociation at RT were used to avoid over digestion and reduce cell membrane damage. This allowed long recording periods (up to 2 h) when necessary. Cells were stored in Eppendorf tube at RT until they were plated. Cells were plated inside cell-culture inserts (IBIDI, Cat# 80209) attached to a 12 mm coverslip (2-4 per day). At DIV2-5, cells were exposed to 500 U/mL hIFN-α either during electrophysiological recordings (acute), or preincubated 24-48 h before recordings.

### ICC and image analysis

Purified recombinant human hIFN-α2a (Sigma-Aldrich, cat. no IF007) or hIFN-β 1a (Sigma-Aldrich, cat. no IF014) were purchased from Millipore. hDRG neuronal dissociated cultures (*DIV* 5-6) were incubated with 300 or 500 U/mL of IFN or vehicle at 37°C with 5% CO_2_. IFNs were applied to hDRG cultures for 30 min and 1 h for immunocytochemistry (ICC) experiments. For the immunodetection of p-eIF4E and p-STAT1 in cultured neurons, hDRG cultures were fixed with 10% formalin (ThermoFisher Scientific, cat. no. 23-245684) at pH 7.4 for 10 min. The chambers were then rinsed three times with 0.1 M PB and blocked with 10% NGS and 0.3% Triton X-100 in 0.1 M PB for 1 h at RT. Next, they were incubated overnight with the primary antibodies anti-p-STAT1 (Tyr701, 1:500; Cell Signaling Technology), or anti-p-eIF4E (1:2000, Abcam cat. no. ab76265), and α-peripherin (1:1000, EnCor Biotechnology, cat. no. cpca-peri). After rinsing three times with 0.1 M PB, the cultures were incubated with secondary antibodies goat anti-chicken IgY (1:2000, H+L, Alexa Fluor 488, Invitrogen, cat. no. A11039) and goat anti-rabbit IgG (1:2000, H+L, Alexa Fluor 647, Thermo Fisher Scientific, cat. no. A21245). The chambers were rinsed 3 times with 0.1 M PB and mounted with ProLong™ Gold Antifade Mountant (Fisher Scientific cat. no. P36930). Slides were examined and imaged at 20X magnification for p-eIF4E and 40X for p-STAT1 immunodetection using an Olympus FV3000 RS confocal laser scanning microscope. Negative controls were performed by the omission of the primary antibody, which resulted in the absence of immunofluorescence. Images were analyzed using OLYMPUS CellSens Dimension software version 1.18.16686.0 by drawing ROIs on individual neuronal cell bodies and acquiring mean gray intensity values for the targeted fluorophores. An extra channel (Alexa Fluor 555) was used to subtract the lipofuscin autofluorescence signal. To obtain cell surface IFNAR1 and IFNAR2 fluorescence profiles, a line (ROI) perpendicular to the cell surface was hand-drawn on ImageJ on each cell culture stained for IFNAR1 and IFNAR2 stimulated with type I IFNs or vehicle. This line started outside the cell and was extended ∼10 microns towards the cell center. Mean gray intensity values along the ROIs were plotted for each protein. To avoid nonspecific signal outside the cells of interest, values below 5% of the maximum mean gray intensity value per cell were considered zero.

### Calcium imaging and analysis

hDRG neurons were cultured on 12-mm glass coverslips coated with 0.1 mg/mL poly-d-lysine hydrobromide (Sigma Aldrich, cat. no. P7405). After pre-treatment with hIFN-α or hIFN-β, cells were washed once with media and loaded with 10 uM Fluo-4 AM (Thermo Scientific, cat. no. F142010) with 0.04% Pluronic™ F-127 (Thermo Scientific, cat. no. P3000MP) in HBSS for 30 min prior to imaging. The coverslip was mounted onto imaging chamber (Warner Instruments, cat. no. RC-25). Cells were washed with recording solution (130 mM NaCl, 4.2 mM KCl, 1.1 mM CaCl_2_, 1 mM MgSO_4_, 0.45 mM NaH_2_PO_4_.H_2_O, 0.5 mM NaH_2_PO_4_.7H_2_O, 20 mM glucose, and 10 mM HEPES, osmolarity 300-310 mOsm and pH 7.4 adjusted with N-methyl-D-glucamine) for 5 min at a flow rate of 1 mL/min followed by 10 min of baseline recording. A high-K^+^ solution was made by adjusting KCl to 50 mM and NaCl to 84.2 mM in the recording solution. Cells that did not fluctuate during baseline recording and exhibited 10% change in response to 200 nM capsaicin were considered for analysis. Calcium responses were captured using an Olympus IX83 inverted microscope at magnification of 10X and MetaFluor for Olympus imaging software (version 7.10.5.476) (71). Elve Flow MUX Distribution 12-way Bidirectional Valve was used to perfuse solutions over the cells. All recordings were performed within DIV 1-4 at RT.

### MEA

Primary human sensory neurons were procured and dissociated as described above. Multi-well microelectrode array culture plates (6 well Cytoview, Cat. M384-tMEA-6W, Axion Biosystems) were prepared by coating the center of the array with 7 µL 0.01% poly-L-ornithine (PLO, EMD Millipore Corp, cat. no. A-004-C). The PLO was left overnight at room temperature. The following day, PLO was removed, and each well was washed three times with sterile DI water and allowed to dry inside the culture cabinet. Once the wells were completely dry, 7 µL laminin (iMatrix silk-511, Matrixome Inc. Cat. no. 892021, at 1:50 in dPBS without calcium or magnesium) were added to the center of the array to ensure that the cells adhered to the area where the electrodes were located. The MEA was maintained at 37°C under 5% CO_2_ for 2 h. Immediately prior to seeding cells, laminin was removed and a 7 µL droplet with approximately 300 neurons was added to the center of the well. Cell attachment was confirmed using a phase-contrast microscope, and the well was flooded with 1 mL of supplemented culture media, as described in the hDRG culture preparation section. The wells were left inside an incubator and maintained with a 50% media change every other day.

At DIV 6, each 6-well MEA was taken to a Maestro Pro acquisition system (Axion Biosystems), and AP waveforms were registered for 15 min under controlled environmental conditions (humidity, 5% CO_2_ and 37 °C) via the enclosed hardware setup. Data were sampled using AxIS software (Axion Biosystems) from raw voltage traces with a bandpass filter from 200 to 3000 Hz. APs were detected from the filtered voltage trace, and individual units were sampled with an adaptive 5.5 SD sliding threshold window in AxIS Navigator Version 3.7.2.8. For the experiment, 15 min were sampled prior to intervention, after which 50% of the media (500 µL) was removed per well and mixed into a final concentration for each assigned group as follows: a) 500U/mL hIFN-α, b) 500 U/mL hIFN-β, c) vehicle for hIFN-α treatment, d) vehicle for hIFN-β treatment, e) hIFNs + eFT508 25 nM, f) vehicle for hIFNs + veh for eFT508. Each group consisted of 4 independent wells derived from duplicate experiments and multiple donors (Table S). At 24 h of treatment, the MEA was registered inside the Maestro Pro system to assess APs after the addition of 100 nM capsaicin to each well.

Files were converted from “.spk” to “.nex” format using the Axion Data Export tool (Axion Biosystems) and analyzed in NeuroExplorer (Plexon, Inc., version 5.425) to assess the two components. First, rate histograms of the evoked spikes were created from a per electrode basis in 50-ms bins and plotted in excel for each group to extract the latency to the first spike after the addition of capsaicin. Then, 1-s bins were used to plot the temporal effect for each group derived from the addition of capsaicin.

### Electrophysiological recordings

Human DRG neurons were recorded 2-7 days after plating (see dissociation details above). The electrophysiological properties of hDRG neurons have been reported to be stable within this range of days *in vitro* (48). The external solution contained (in mM): 125 NaCl, 3.6 KCl, 2 NaH_2_PO_4_, 2.5 CaCl_2_, 2 MgCl_2_, 25 NaHCO_3_, and 11 dextrose. Whole-cell patch clamp was performed using borosilicate capillaries pulled with a P-97 flaming-brown micropipette puller (Sutter Instruments). The pipettes had a resistance of 2-4 MΩ, when using an internal solution containing (in mM): 120 K-Glutamate, 2 KCl, 8 NaCl, 0.2 EGTA, 14 Na2-Phoshphocreatine, 2 Mg-ATP, 0.3 Na-GTP and 10 HEPES-K (pH 7.3, 295 mOsm). Recordings were obtained using an Axopatch 200B amplifier (Molecular Devices). Neurons were visualized using a Nikon Eclipse Ti inverted microscope equipped with Nikon Advanced Modulation Contrast. Acquisition was done at 10-20 kHz and data was filtered at 5 kHz and analyzed offline using Clampfit Analysis Suite 11 (Molecular Devices), GraphPad Prism 11, and Microsoft Excel 2015.

After achieving whole-cell configuration, cells were held near resting potential (Vm_rest_) for at least 5 min to allow for dialysis of the pipette internal solution. Afterwards, when needed, cells were held at -70 mV by injecting current of the appropriate sign and amplitude. Depending on the experiment, one or several of the following protocols were applied: For rheobase evaluation, first, a gross determination of action potential firing was made using 10-200 pA incremental steps (500 ms duration), and then a baseline subthreshold current with incremental steps of 1/10th of the apparent Rh in successive sweeps were used. The rheobase was considered the step where the first AP was fired and where APs were consistently fired with repeat step stimulation. The maximum stimulation intensity used to test rheobase was 18 nA. This maximum value was considered the rheobase for cells that did not fire action potentials. To further determine the excitability characteristics of each cell, they were subjected to 12 incremental steps of magnitude 1/5th that of a baseline set at 1.2X the rheobase. For acute exposure to hIFN-α, 10-15 baseline sweeps were recorded. Then, 500U/mL of hIFN-α were added and recordings continued for 30-90 min.

### Statistics

Data were plotted and analyzed using GraphPad Prism version 10.0.2232 (San Diego, CA). All data are shown as mean ± standard error of the mean (SEM). Student’s t-test, one-way or two-way ANOVA followed by Bonferroni test were used to analyze data. The test used is indicated for each experiment in the results section.

## Author Contributions

UFE and TJP conceived the idea and designed research project and experiments; UFE and KN performed IHC and in situ hybridization, FE performed electrophysiological recordings, RGV performed microelectrode recordings; KN performed calcium imaging; UFE, KN, FE, RGV and HM analyzed data; UFE and TJP wrote the paper. All authors edited the paper and provided input to the final manuscript.

## Acknowledgements

The authors thank Erin Vines, Anna Cervantes, Geoffrey Funk, and Peter Horton at the Southwest Transplant Alliance. The authors are grateful to the organ donors and their families for their gift.

## References

1. Garcia-Sastre, A., and Biron, C.A. 2006. Type 1 interferons and the virus-host relationship: a lesson in detente. Science 312:879–882.

2. McNab, F., Mayer-Barber, K., Sher, A., Wack, A., and O’Garra, A. 2015. Type I interferons in infectious disease. Nat Rev Immunol 15:87–103.

3. Karakoese, Z., Ingola, M., Sitek, B., Dittmer, U., and Sutter, K. 2024. IFNalpha Subtypes in HIV Infection and Immunity. Viruses 16.

4. Zanin, N., Viaris de Lesegno, C., Lamaze, C., and Blouin, C.M. 2020. Interferon Receptor Trafficking and Signaling: Journey to the Cross Roads. Front Immunol 11:615603.

5. Piehler, J., Thomas, C., Garcia, K.C., and Schreiber, G. 2012. Structural and dynamic determinants of type I interferon receptor assembly and their functional interpretation. Immunol Rev 250:317–334.

6. Shemesh, M., Lochte, S., Piehler, J., and Schreiber, G. 2021. IFNAR1 and IFNAR2 play distinct roles in initiating type I interferon-induced JAK-STAT signaling and activating STATs. Sci Signal 14:eabe4627.

7. Boxx, G.M., and Cheng, G. 2016. The Roles of Type I Interferon in Bacterial Infection. Cell Host Microbe 19:760–769.

8. Mazewski, C., Perez, R.E., Fish, E.N., and Platanias, L.C. 2020. Type I Interferon (IFN)-Regulated Activation of Canonical and Non-Canonical Signaling Pathways. Front Immunol 11:606456.

9. Pujantell, M., and Altfeld, M. 2022. Consequences of sex differences in Type I IFN responses for the regulation of antiviral immunity. Front Immunol 13:986840.

10. Fitzgibbon, M., Kerr, D.M., Henry, R.J., Finn, D.P., and Roche, M. 2019. Endocannabinoid modulation of inflammatory hyperalgesia in the IFN-alpha mouse model of depression. Brain Behav Immun 82:372–381.

11. Barragan-Iglesias, P., Franco-Enzastiga, U., Jeevakumar, V., Shiers, S., Wangzhou, A., Granados-Soto, V., Campbell, Z.T., Dussor, G., and Price, T.J. 2020. Type I Interferons Act Directly on Nociceptors to Produce Pain Sensitization: Implications for Viral Infection-Induced Pain. J Neurosci 40:3517–3532.

12. Franco-Enzastiga, U., Natarajan, K., David, E.T., Patel, K., Ravirala, A., and Price, T.J. 2024. Vinorelbine causes a neuropathic pain-like state in mice via STING and MNK1 signaling associated with type I interferon induction. iScience 27:108808.

13. Lin, C.Y., Guu, T.W., Lai, H.C., Peng, C.Y., Chiang, J.Y., Chen, H.T., Li, T.C., Yang, S.Y., Su, K.P., and Chang, J.P. 2020. Somatic pain associated with initiation of interferon-alpha (IFN-alpha) plus ribavirin (RBV) therapy in chronic HCV patients: A prospective study. Brain Behav Immun Health 2:100035.

14. Ray, P.R., Shiers, S., Caruso, J.P., Tavares-Ferreira, D., Sankaranarayanan, I., Uhelski, M.L., Li, Y., North, R.Y., Tatsui, C., Dussor, G., et al. 2023. RNA profiling of human dorsal root ganglia reveals sex differences in mechanisms promoting neuropathic pain. Brain 146:749–766.

15. Li, Y., Gray, E.H., Ross, R., Zebochin, I., Lock, A., Fedele, L., Kamajaya, L.J., Marrow, R.J., Ryan, S., Röderer, P., et al. 2024. Blockade of rheumatoid arthritis synovial fluid-induced sensory neuron activation by JAK inhibitors. bioRxiv:2024.2008.2019.608613.

16. Crow, M.K. 2014. Type I interferon in the pathogenesis of lupus. J Immunol 192:5459–5468.

17. Su, J., Zhang, M.-D., Kupari, J., Kwak, D., Picton, L., Xu, B., Hu, Y., Alvarez, A.G., Usoskin, D., Xu, Z., et al. 2025. Persistent interferon signaling that causes sensory neuron plasticity and pain in arthritis. bioRxiv:2025.2001.2018.633447.

18. Li, Y., Uhelski, M.L., North, R.Y., Mwirigi, J.M., Tatsui, C.E., McDonough, K.E., Cata, J.P., Corrales, G., Dussor, G., Price, T.J., et al. 2024. Tomivosertib reduces ectopic activity in dorsal root ganglion neurons from patients with radiculopathy. Brain 147:2991–2997.

19. Shiers, S.I., Mazhar, K., Wangzhou, A., Haberberger, R., Lesnak, J.B., Sankaranarayanan, I., Tavares-Ferreira, D., Cervantes, A., Funk, G., Horton, P., et al. 2024. Nageotte nodules in human DRG reveal neurodegeneration in painful diabetic neuropathy. bioRxiv.

20. Herdy, B., Jaramillo, M., Svitkin, Y.V., Rosenfeld, A.B., Kobayashi, M., Walsh, D., Alain, T., Sean, P., Robichaud, N., Topisirovic, I., et al. 2012. Translational control of the activation of transcription factor NF-kappaB and production of type I interferon by phosphorylation of the translation factor eIF4E. Nat Immunol 13:543–550.

21. Moy, J.K., Kuhn, J.L., Szabo-Pardi, T.A., Pradhan, G., and Price, T.J. 2018. eIF4E phosphorylation regulates ongoing pain, independently of inflammation, and hyperalgesic priming in the mouse CFA model. Neurobiol Pain 4:45–50.

22. Moy, J.K., Khoutorsky, A., Asiedu, M.N., Black, B.J., Kuhn, J.L., Barragan-Iglesias, P., Megat, S., Burton, M.D., Burgos-Vega, C.C., Melemedjian, O.K., et al. 2017. The MNK-eIF4E Signaling Axis Contributes to Injury-Induced Nociceptive Plasticity and the Development of Chronic Pain. J Neurosci 37:7481–7499.

23. Wong, C., Tavares-Ferreira, D., Thorn Perez, C., Sharif, B., Uttam, S., Amiri, M., Lister, K.C., Hooshmandi, M., Nguyen, V., Seguela, P., et al. 2023. 4E-BP1-dependent translation in nociceptors controls mechanical hypersensitivity via TRIM32/type I interferon signaling. Sci Adv 9:eadh9603.

24. Megat, S., Ray, P.R., Moy, J.K., Lou, T.F., Barragan-Iglesias, P., Li, Y., Pradhan, G., Wanghzou, A., Ahmad, A., Burton, M.D., et al. 2019. Nociceptor Translational Profiling Reveals the Ragulator-Rag GTPase Complex as a Critical Generator of Neuropathic Pain. J Neurosci 39:393–411.

25. Shiers, S., Mwirigi, J., Pradhan, G., Kume, M., Black, B., Barragan-Iglesias, P., Moy, J.K., Dussor, G., Pancrazio, J.J., Kroener, S., et al. 2020. Reversal of peripheral nerve injury-induced neuropathic pain and cognitive dysfunction via genetic and tomivosertib targeting of MNK. Neuropsychopharmacology 45:524–533.

26. Lackovic, J., Price, T.J., and Dussor, G. 2023. MNK1/2 contributes to periorbital hypersensitivity and hyperalgesic priming in preclinical migraine models. Brain 146:448–454.

27. David, E.T., Yousuf, M.S., Mei, H.R., Jain, A., Krishnagiri, S., Elahi, H., Venkatesan, R., Srikanth, K.D., Dussor, G., Dalva, M.B., et al. 2024. ephrin-B2 promotes nociceptive plasticity and hyperalgesic priming through EphB2-MNK-eIF4E signaling in both mice and humans. Pharmacol Res 206:107284.

28. Mitchell, M.E., Torrijos, G., Cook, L.F., Mwirigi, J.M., He, L., Shiers, S., and Price, T.J. 2024. Interleukin-6 induces nascent protein synthesis in human dorsal root ganglion nociceptors primarily via MNK-eIF4E signaling. Neurobiol Pain 16:100159.

29. Defaye, M., Bradaia, A., Abdullah, N.S., Agosti, F., Iftinca, M., Delanne-Cumenal, M., Soubeyre, V., Svendsen, K., Gill, G., Ozmaeian, A., et al. 2024. Induction of antiviral interferon-stimulated genes by neuronal STING promotes the resolution of pain in mice. J Clin Invest 134.

30. Donnelly, C.R., Jiang, C., Andriessen, A.S., Wang, K., Wang, Z., Ding, H., Zhao, J., Luo, X., Lee, M.S., Lei, Y.L., et al. 2021. STING controls nociception via type I interferon signalling in sensory neurons. Nature 591:275–280.

31. Wang, K., Donnelly, C.R., Jiang, C., Liao, Y., Luo, X., Tao, X., Bang, S., McGinnis, A., Lee, M., Hilton, M.J., et al. 2021. STING suppresses bone cancer pain via immune and neuronal modulation. Nat Commun 12:4558.

32. Silveira Prudente, A., Hoon Lee, S., Roh, J., Luckemeyer, D.D., Cohen, C.F., Pertin, M., Park, C.K., Suter, M.R., Decosterd, I., Zhang, J.M., et al. 2024. Microglial STING activation alleviates nerve injury-induced neuropathic pain in male but not female mice. Brain Behav Immun 117:51–65.

33. Liu, C.C., Gao, Y.J., Luo, H., Berta, T., Xu, Z.Z., Ji, R.R., and Tan, P.H. 2016. Interferon alpha inhibits spinal cord synaptic and nociceptive transmission via neuronal-glial interactions. Sci Rep 6:34356.

34. Liu, S., Karaganis, S., Mo, R.F., Li, X.X., Wen, R.X., and Song, X.J. 2020. IFNbeta Treatment Inhibits Nerve Injury-induced Mechanical Allodynia and MAPK Signaling By Activating ISG15 in Mouse Spinal Cord. J Pain 21:836–847.

35. Tan, P.H., Gao, Y.J., Berta, T., Xu, Z.Z., and Ji, R.R. 2012. Short small-interfering RNAs produce interferon-alpha-mediated analgesia. Br J Anaesth 108:662–669.

36. Woller, S.A., Ocheltree, C., Wong, S.Y., Bui, A., Fujita, Y., Goncalves Dos Santos, G., Yaksh, T.L., and Corr, M. 2019. Neuraxial TNF and IFN-beta co-modulate persistent allodynia in arthritic mice. Brain Behav Immun 76:151–158.

37. Kalie, E., Jaitin, D.A., Podoplelova, Y., Piehler, J., and Schreiber, G. 2008. The stability of the ternary interferon-receptor complex rather than the affinity to the individual subunits dictates differential biological activities. J Biol Chem 283:32925–32936.

38. Schreiber, G. 2017. The molecular basis for differential type I interferon signaling. J Biol Chem 292:7285–7294.

39. Wittling, M.C., Cahalan, S.R., Levenson, E.A., and Rabin, R.L. 2020. Shared and Unique Features of Human Interferon-Beta and Interferon-Alpha Subtypes. Front Immunol 11:605673.

40. Bhuiyan, S.A., Xu, M., Yang, L., Semizoglou, E., Bhatia, P., Pantaleo, K.I., Tochitsky, I., Jain, A., Erdogan, B., Blair, S., et al. 2024. Harmonized cross-species cell atlases of trigeminal and dorsal root ganglia. Science Advances 10:eadj9173.

41. Tavares-Ferreira, D., Shiers, S., Ray, P.R., Wangzhou, A., Jeevakumar, V., Sankaranarayanan, I., Cervantes, A.M., Reese, J.C., Chamessian, A., Copits, B.A., et al. 2022. Spatial transcriptomics of dorsal root ganglia identifies molecular signatures of human nociceptors. Sci Transl Med 14:eabj8186.

42. Hadjilambreva, G., Mix, E., Rolfs, A., Muller, J., and Strauss, U. 2005. Neuromodulation by a cytokine: interferon-beta differentially augments neocortical neuronal activity and excitability. J Neurophysiol 93:843–852.

43. Hallal, D.E., Farias, A.S., Oliveira, E.C., Diaz-Bardales, B.M., Brandao, C.O., Protti, G.G., Pereira, F.G., Metze, I.L., and Santos, L.M. 2003. Costimulatory molecule expression on leukocytes from mice with experimental autoimmune encephalomyelitis treated with IFN-beta. J Interferon Cytokine Res 23:293–298.

44. Jaitin, D.A., Roisman, L.C., Jaks, E., Gavutis, M., Piehler, J., Van der Heyden, J., Uze, G., and Schreiber, G. 2006. Inquiring into the differential action of interferons (IFNs): an IFN-alpha2 mutant with enhanced affinity to IFNAR1 is functionally similar to IFN-beta. Mol Cell Biol 26:1888–1897.

45. Holder, P.G., Lim, S.A., Huang, C.S., Sharma, P., Dagdas, Y.S., Bulutoglu, B., and Sockolosky, J.T. 2022. Engineering interferons and interleukins for cancer immunotherapy. Adv Drug Deliv Rev 182:114112.

46. Marijanovic, Z., Ragimbeau, J., van der Heyden, J., Uze, G., and Pellegrini, S. 2007. Comparable potency of IFNalpha2 and IFNbeta on immediate JAK/STAT activation but differential down-regulation of IFNAR2. Biochem J 407:141–151.

47. Tolomeo, M., Cavalli, A., and Cascio, A. 2022. STAT1 and Its Crucial Role in the Control of Viral Infections. Int J Mol Sci 23.

48. Davidson, S., Copits, B.A., Zhang, J., Page, G., Ghetti, A., and Gereau, R.W.t. 2014. Human sensory neurons: Membrane properties and sensitization by inflammatory mediators. Pain 155:1861–1870.

49. Zurek, N.A., Ehsanian, R., Goins, A.E., Adams, I.M., Petersen, T., Goyal, S., Shilling, M., Westlund, K.N., and Alles, S.R.A. 2024. Electrophysiological Analyses of Human Dorsal Root Ganglia and Human Induced Pluripotent Stem Cell-derived Sensory Neurons From Male and Female Donors. J Pain 25:104451.

50. Sego, T.J., Aponte-Serrano, J.O., Ferrari Gianlupi, J., Heaps, S.R., Breithaupt, K., Brusch, L., Crawshaw, J., Osborne, J.M., Quardokus, E.M., Plemper, R.K., et al. 2020. A modular framework for multiscale, multicellular, spatiotemporal modeling of acute primary viral infection and immune response in epithelial tissues and its application to drug therapy timing and effectiveness. PLoS Comput Biol 16:e1008451.

51. Yi, J., Yang, L., Widman, A.J., Toliver, A., Bertels, Z., Del Rosario, J.S., Slivicki, R.A., Payne, M., Dourson, A.J., Li, J.-N., et al. 2025. Human sensory neurons exhibit cell-type-specific, pain-associated differences in intrinsic excitability and expression of SCN9A and SCN10A bioRxiv:2025.2003.2025.645367.

52. Waskiewicz, A.J., Johnson, J.C., Penn, B., Mahalingam, M., Kimball, S.R., and Cooper, J.A. 1999. Phosphorylation of the cap-binding protein eukaryotic translation initiation factor 4E by protein kinase Mnk1 in vivo. Mol Cell Biol 19:1871–1880.

53. Kalia, A.K., Rosseler, C., Granja-Vazquez, R., Ahmad, A., Pancrazio, J.J., Neureiter, A., Zhang, M., Sauter, D., Vetter, I., Andersson, A., et al. 2023. How to differentiate induced pluripotent stem cells into sensory neurons for disease modelling: a comparison of two protocols. Res Sq.

54. Lee, S.H., Bonifacio, F., Prudente, A.S., Choi, Y.I., Roh, J., Adjafre, B.L., Park, C.K., Jung, S.J., Cunha, T.M., and Berta, T. 2024. STING recognition of viral dsDNA by nociceptors mediates pain in mice. Brain Behav Immun 121:29–42.

55. Lei, V., Handfield, C., Kwock, J.T., Kirchner, S.J., Lee, M.J., Coates, M., Wang, K., Han, Q., Wang, Z., Powers, J.G., et al. 2022. Skin Injury Activates a Rapid TRPV1-Dependent Antiviral Protein Response. J Invest Dermatol 142:2249–2259 e2249.

56. Sen, N., Sung, P., Panda, A., and Arvin, A.M. 2018. Distinctive Roles for Type I and Type II Interferons and Interferon Regulatory Factors in the Host Cell Defense against Varicella-Zoster Virus. J Virol 92.

57. Sun, Y., Wang, G., Wang, R., Ren, L., Yuan, Z., Liu, Y., Wu, Y., Chen, R., Chen, Y., and Diao, B. 2023. Serum levels of type I interferon (IFN-I) is associated with the severity of COVID-19. J Med Microbiol 72.

58. Zhang, H., Li, A., Liu, Y.F., Sun, Z.M., Jin, B.X., Lin, J.P., Yang, Y., and Yao, Y.X. 2023. Spinal TAOK2 contributes to neuropathic pain via cGAS-STING activation in rats. iScience 26:107792.

59. Reich, S.H., Sprengeler, P.A., Chiang, G.G., Appleman, J.R., Chen, J., Clarine, J., Eam, B., Ernst, J.T., Han, Q., Goel, V.K., et al. 2018. Structure-based Design of Pyridone-aminal eFT508 Targeting Dysregulated Translation by Selective Mitogen-activated Protein Kinase Interacting Kinases 1 and 2 (MNK1/2) Inhibition. J Med Chem.

60. Moy, J.K., Khoutorsky, A., Asiedu, M.N., Dussor, G., and Price, T.J. 2018. eIF4E Phosphorylation Influences Bdnf mRNA Translation in Mouse Dorsal Root Ganglion Neurons. Front Cell Neurosci 12:29.

61. Preston, S.E.J., Dahabieh, M.S., Flores Gonzalez, R.E., Goncalves, C., Richard, V.R., Leibovitch, M., Dakin, E., Papadopoulos, T., Lopez Naranjo, C., McCallum, P.A., et al. 2024. Blocking tumor-intrinsic MNK1 kinase restricts metabolic adaptation and diminishes liver metastasis. Sci Adv 10:eadi7673.

62. Walsh, D., and Mohr, I. 2004. Phosphorylation of eIF4E by Mnk-1 enhances HSV-1 translation and replication in quiescent cells. Genes Dev 18:660–672.

63. Paul, F., Pellegrini, S., and Uze, G. 2015. IFNA2: The prototypic human alpha interferon. Gene 567:132–137.

64. Moreau, T.R.J., Bondet, V., Rodero, M.P., and Duffy, D. 2023. Heterogeneity and functions of the 13 IFN-alpha subtypes - lucky for some? Eur J Immunol 53:e2250307.

65. Patil, M.J., Ru, F., Sun, H., Wang, J., Kolbeck, R.R., Dong, X., Kollarik, M., Canning, B.J., and Undem, B.J. 2020. Acute activation of bronchopulmonary vagal nociceptors by type I interferons. J Physiol 598:5541–5554.

66. He, D., Mao, A., Li, Y., Tam, S., Zheng, Y., Yao, X., Birnbaumer, L., Ambudkar, I.S., and Ma, X. 2020. TRPC1 participates in the HSV-1 infection process by facilitating viral entry. Sci Adv 6:eaaz3367.

67. Guo, Y.Y., Gao, Y., Zhao, Y.L., Xie, C., Gan, H., Cheng, X., Yang, L.P., Hu, J., Shu, H.B., Zhong, B., et al. 2024. Viral infection and spread are inhibited by the polyubiquitination and downregulation of TRPV2 channel by the interferon-stimulated gene TRIM21. Cell Rep 43:114095.

68. Buttari, F., Zagaglia, S., Marciano, L., Albanese, M., Landi, D., Nicoletti, C.G., Mercuri, N.B., Silvestrini, M., Provinciali, L., Marfia, G.A., et al. 2017. TRPV1 polymorphisms and risk of interferon beta-induced flu-like syndrome in patients with relapsing-remitting multiple sclerosis. J Neuroimmunol 305:172–174.

69. Shiers, S., Yousuf, M.S., Mwirigi, J.M., Cervantes, A., and Price, T.J. 2024. Human Ganglia and Spinal Cord Tissue Procurement from Organ Donors and Tissue Quality Assessment. Protocols.io.

70. Valtcheva, M.V., Copits, B.A., Davidson, S., Sheahan, T.D., Pullen, M.Y., McCall, J.G., Dikranian, K., and Gereau, R.W.t. 2016. Surgical extraction of human dorsal root ganglia from organ donors and preparation of primary sensory neuron cultures. Nat Protoc 11:1877–1888.

71. Natarajan, K., Franco-Enzástiga, U., Espinosa, F., and Price, T. 2024. Calcium imaging in human dorsal root ganglia neurons. protocols.io.

