## Supplementary Fig and Tables for "Type I interferons enhance human dorsal root ganglion nociceptor excitability and induce TRPV1 sensitization"

+1 972-883-4311

Úrzula Franco-Enzástiga

800 W Campbell Rd

BSB10.657

Richardson TX 75080

USA

+1 469-920-0105

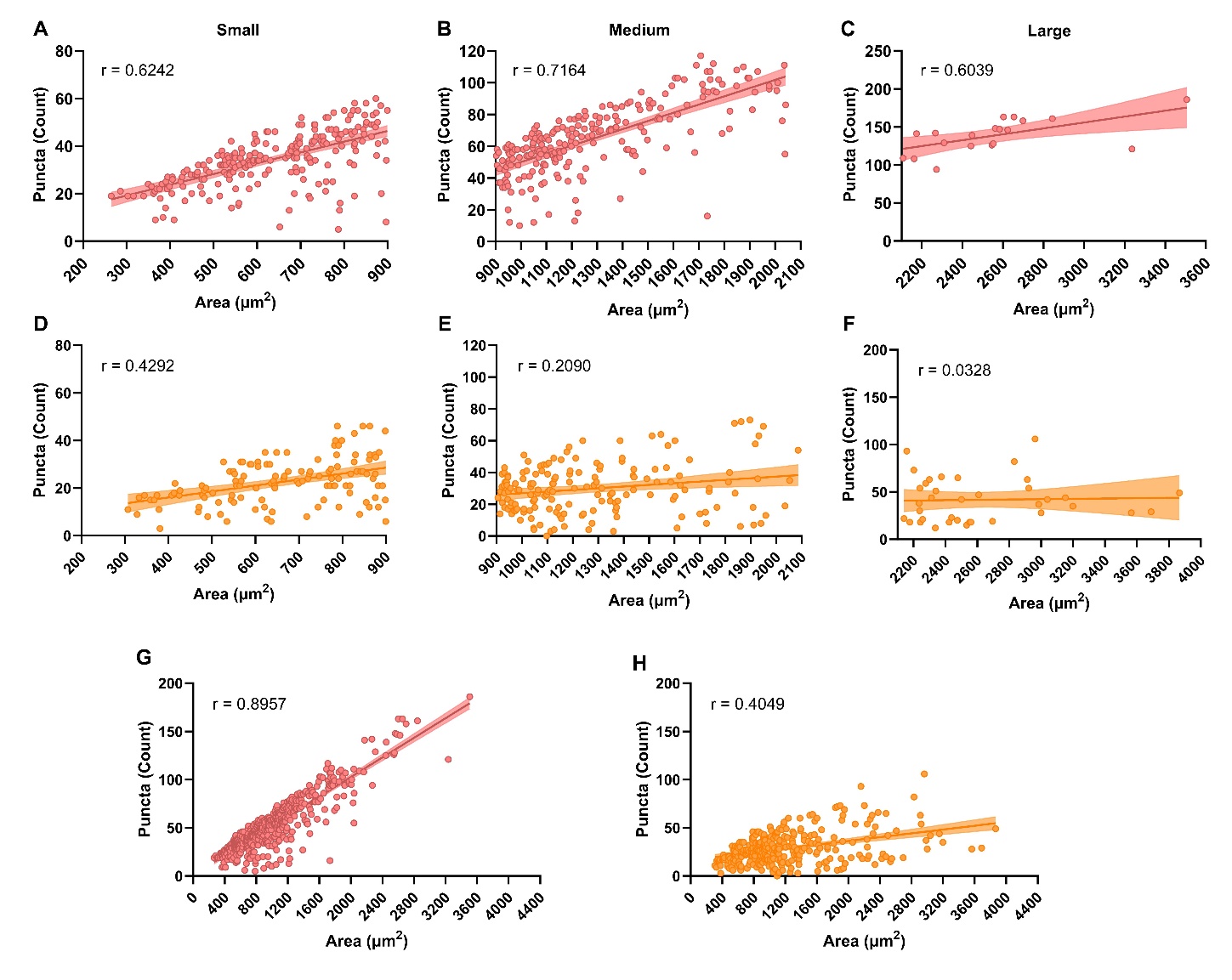
**Fig. S1.** Correlation between number of *IFNAR1* and *IFNAR2* mRNA puncta and hDRG neuronal cell area. **A-C**. Correlation between *IFNAR1* mRNA expression and small **(A)**, medium **(B)**, and large **(C)** cell sizes. **D-F.** Correlation between *IFNAR2* mRNA expression and small **(D)**, medium **(E)**, and large **(F)** cell sizes. **G.** Correlation between *IFNAR1* mRNA expression and all neuronal cell sizes. **H.** Correlation between *IFNAR2* mRNA expression and all neuronal cell sizes.

**A**

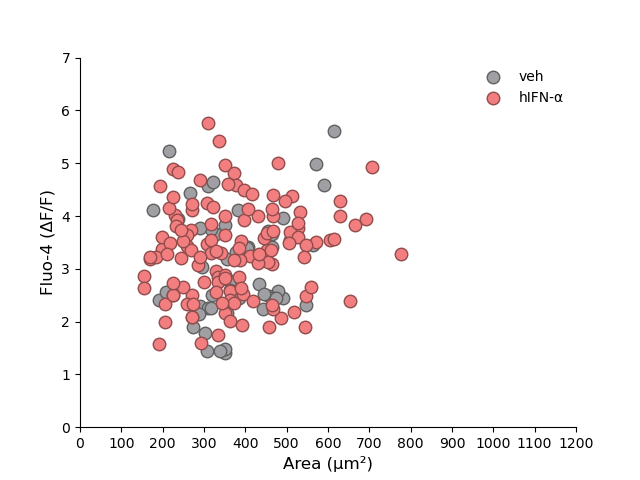

**B**

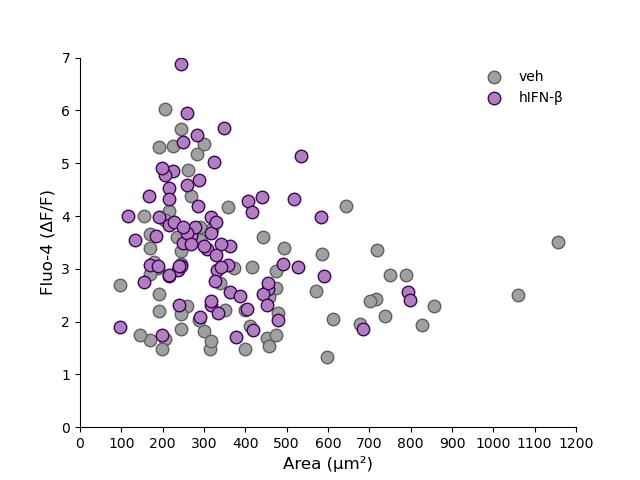

**Fig. S2.** Peak calcium response to capsaicin in hDRG neurons treated with hIFN-α or hIFN-β across different cell sizes. **A.** Peak fold-change in calcium response to capsaicin of hDRG neurons treated with vehicle or hIFN-α plotted as a function of cell area. **B.** Peak fold-change in calcium response to capsaicin of hDRG neurons treated with vehicle or hIFN-β plotted as a function of cell area.

| **Assay** | **Donor** | **Age** | **Sex** | **COD** |
| --- | --- | --- | --- | --- |
| RNAScope | 1 | 29 | F | Anoxia/Drug Overdose |
|  | 2 | 19 | M | Anoxia/Drug Overdose |
|  | 3 | 30 | M | Anoxia/Asthma Attack |
| Treatments and ICCs | 4 | 18 | M | Anoxia/Natural Causes |
|  | 5 | 52 | M | CVA/Stroke |
|  | 6 | 22 | M | Head Trauma/Blunt Injury/MVA |
|  | 7 | 37 | F | Head Trauma/GSW/Homicide |
|  | 8 | 49 | M | CVA/Stroke |
|  | 9* | 29 | M | Head Trauma/Blunt Injury/MVA |
|  | 10 | 27 | M | CVA/Stroke |
|  | 11 | 35 | M | Anoxia/Drug Intoxication |
|  | 12 | 50 | M | CVA/Stroke |
| Calcium Imaging | 13 | 32 | M | Head Trauma/MVA |
|  | 14 | 44 | F | Anoxia/Cardiovascular |
|  | 15 | 33 | M | Head Trauma |
|  | 16 | 46 | M | Anoxia |
|  | 17* | 58 | F | Cardiac Arrest |
| MEA | 18* | 28 | M | Anoxia |
|  | 19* | 49 | M | Head Trauma |
|  | 20* | 18 | M | Head Trauma/GSW |
| Patch-clamp | 21 | 45 | M | Anoxia |
|  | 22 | 34 | F | Anoxia/Drug Overdose |
|  | 23 | 27 | M | CVA/Stroke |
|  | 24 | 29 | F | Head Trauma/GSW/Suicide |
|  | 25 | 11 | M | Anoxia |
|  | 26 | 20 | M | Head Trauma/Blunt Injury/MVA |
|  | 27 | 19 | M | Anoxia/Cardiovascular |
|  | 28 | 32 | M | Anoxia/Drug Intoxication |
|  | 29 | 26 | M | Anoxia/Drug Overdose |
|  | 30 | 26 | M | Anoxia |
|  | 31 | 58 | F | Anoxia |
|  | 32 | 50 | M | Anoxia |
|  | 33 | 25 | M | Anoxia Secondary to MI |
| * Indicates donors also used for patch-clamp experiments. | | | | |

**Table S1** Donor demographic data
